# Peptide Coated Polycaprolactone-Benzalkonium Chloride Nanocapsules for Targeted Drug Delivery to the Pancreatic β-Cell

**DOI:** 10.1101/2024.07.15.603612

**Authors:** Jillian Collins, Jessie M. Barra, Kiefer Holcomb, Andres Ocampo, Ashton Fremin, Jubril Akolade, Austin Kratz, Julianna K. Hays, Ali Shilleh, David J. Hodson, Johannes Broichhagen, Holger A. Russ, Nikki L. Farnsworth

## Abstract

Targeting of current therapies to treat or prevent loss of pancreatic islet β-cells in Type 1 Diabetes (T1D) may provide improved efficacy and reduce off target effects. Current efforts to target the β-cell are limited by a lack of β-cell specific targets and the inability to test multiple targeting moieties with the same delivery vehicle. Here we fabricate a novel tailorable polycaprolactone nanocapsule (NC) where multiple different targeting peptides can be interchangeably attached for β-cell specific delivery. Incorporation of a cationic surfactant in the NC shell allows for the attachment of Exendin-4 and an antibody for ectonucleoside triphosphate diphosphohydrolase 3 (ENTPD3) for β-cell specific targeting. The average NC size ranges from 250-300nm with a polydispersity index under 0.2. The NCs are non-toxic, stable in media culture, and can be lyophilized and reconstituted. NCs coated with targeting peptide were taken up by human cadaveric islet β-cells and human stem cell-derived β-like cells (sBC) *in vitro* with a high level of specificity. Furthermore, NCs successfully delivered both hydrophobic and hydrophilic cargo to human β-cells. Finally, Exendin-4 coated NCs were stable and targeted the mouse pancreatic islet β-cell *in vivo*. Our unique NC design allows for the interchangeable coating of targeting peptides for future screening of targets with improved cell specificity. The ability to target and deliver therapeutics to human pancreatic β-cells opens avenues for improved therapies and treatments to help the delay onset, prevent, or reverse T1D.

---

Type 1 diabetes (T1D) is a global health problem affecting 8.4 million people worldwide and the number of diagnosed cases is projected to double by 2040.^1^ T1D is characterized by the immune-mediated destruction of insulin producing pancreatic islet β-cells.^1^ This leads to impaired glucose homeostasis and a significant decline in quality of life.^2^ Although exogenous insulin injections can restore glucose homeostasis; improved therapies are needed to decrease the risk of life-threatening disease complications and stop disease progression.^3^ There are a variety of therapies in development to protect β-cell mass and function as a preventative treatment for T1D.^4^ Developing pharmacologic approaches to promote proliferation of residual β-cells or preserve graft mass and function post islet transplantation therapy could represent an alternative to the gold standard insulin injection treatment for established T1D.^4^ Dual specificity tyrosine-phosphorylation-regulated kinase 1A (DYRK1A) inhibitors for example are a class of molecules that can induce proliferation of human β-cells.^5^ However, DYRK1A inhibitors can also inhibit proliferation at higher doses, have significant off-target effects on the nervous system limiting therapeutic efficacy and are not specifically targeted to the β-cell.^5,6^ Other therapeutics in development include the calcium channel blocker Verapamil, which promotes β-cell function in adults with new-onset T1D; however, targeted delivery of increased concentrations of Verapamil to the β-cell may further improve β-cell function and survival without off-target effects of hypotension that occur with oral administration.^7^ Therefore, there is a critical need for targeted drug delivery specifically to the human β-cell for treating T1D.

Intravenous administration of therapeutics with effects on the pancreatic β-cell can bypass the gastrointestinal tract, requires lower dosage, and can reach the exocrine pancreas via the vascular system.^8,9^ However, intravenous drug delivery strategies face a myriad of biological barriers before reaching their target site and have significant off target effects. To circumvent these challenges, utilizing nanocarriers for targeted drug delivery has been expanding due their small size and customizable surface chemistry allowing the attachment of targeting moieties to improve cell specificity and overcome biological barriers.^10– 12^Nanocapsules (NCs), have demonstrated their ability to efficiently deliver small peptides and other therapeutics directly to desired cellular targets in a controlled manner.^13,14^ NCs are a type of nanoparticle consisting of an outer biodegradable polymer shell encompassing a liquid or solid core have demonstrated longer release of cargo compared to nanospheres^15,16^ Poly(ε-caprolactone) (PCL) is a hydrophobic semicrystalline polymer extensively used in nanoparticle formulations for drug delivery.^17,18^ In addition to being biocompatible and biodegradable, PCL nanoparticles have exhibited slow and sustained release of a variety of cargo including small molecules, peptides, and RNA.^19–21^ Furthermore, PCL NCs composed of an oil core encapsulated inside a PCL polymer shell can successfully encapsulate hydrophilic or hydrophobic cargo and deliver therapeutics in a controlled manner.^15,22^ Although nanoparticles provide controlled drug release, cell specific targeting remains challenging due to a lack of cell-specific surface markers.^12,23,24^ This supports a pressing need for innovative drug delivery platforms that provide a higher throughput screening method for cell-specificity with multiple cell surface targets.^11,25^

Nanoparticles targeted to the pancreatic islet β-cell have previously been utilized as a non-invasive strategy for quantifying and visualizing β-cell mass.^26,27^ Notably, iron nanoparticles coated with Exendin-4 (Ex4), an agonist for the glucagon-like peptide 1 receptor (GLP-1R) that is expressed on the surface of β-cells, was successful for targeting the pancreatic islet *in vivo*.^26,28,29^ Although these magnetic nanoparticles could potentially be used as an early diagnostic tool for diabetes detection,^30,31^ implementing nanoparticles for targeted drug delivery to protect and promote the proliferation of pancreatic β-cells has not been fully investigated. The GLP-1R is an attractive target for β-cell specific drug delivery as it’s expression is specific to the β-cell within the islet, the FDA-approved agonist Ex4 exhibits high potency for the GLP-1R, and the internalization of GLP-1R upon ligand binding in pancreatic β-cells has been well characterized.^32,33^ However, while GLP-1R has been used in several studies to target the islet β-cell for imaging or delivery of therapeutics, accumulation of Ex4 targeted cargo in off-target tissues such as the spleen, kidney, liver, and lungs has been observed, and may be attributed to GLP-1R expression in the lungs and liver.^33,34^ Therefore, to improve targeted delivery to islet β-cells, a tailorable drug delivery vehicle with interchangeable targeting moieties is needed to identify novel peptides with high β-cell specificity.

In this study we synthesized a novel NC drug delivery platform, using PCL NCs coated with the cationic surfactant benzalkonium chloride (BKC)-PCL-NCs. The charged surfactant allows adherence of different moieties for specific cell targeting. We demonstrated the NCs can target pancreatic β-cells using two different targeting molecules on the NC surface highlighting the modular capabilities of our approach. Notably, NCs can encapsulate either hydro- or hydrophilic cargo with sustained release for prolonged periods (∼40 days) and are stable under physiological conditions. NCs could be coated with either Ex4 or an antibody for ectonucleoside triphosphate diphosphohydrolase 3 (ENTPD3) for β-cell specific targeting. NCs selectively target human stem cell-derived β-like cells (sBC) and human cadaveric islet β-cells *in vitro* and can deliver therapeutic cargo. Additionally, NCs are stable after tail vein injection into non-obese diabetic/severe combined immunodeficiency (NOD-Scid) mice and can selectively target pancreatic mouse β-cells *in vivo*. The results from this study support proof of principle of targeted delivery to the human β-cell using our innovative NC design. The ability to selectively target and deliver therapeutic cargo to human pancreatic β-cells via NCs significantly improves the prospect of protecting, proliferating, and regenerating replacement, or residual pancreatic β-cells for T1D patients.

## RESULTS AND DISCUSSION

### Feasible to Fabricate BKC-PCL-NCs and Encapsulate Either Hydrophobic or Hydrophilic Cargo

Coated PCL nanoparticles have been utilized in a myriad of drug delivery applications.^15,35,36^ It has been previously demonstrated BKC can integrate into the PCL polymer framework via electrostatic interactions to form solid nanospheres, but has not been demonstrated in a nanocapsule (NC).^37^ Furthermore, BKC’s cationic nature allows the opportunity to adhere targeting peptides to the outside of such particles. BKC is an FDA approved preservative already used for ophthalmic applications and is capable of coating nanoparticles for clinical applications.^36,38,39^ Therefore, we modified existing NP protocols and optimized them to achieve stable NCs with BKC incorporated into the PCL shell (BKC-PCL NCs).^37,40^ For hydrophobic cargo, NCs were synthesized via water in oil (W/O) emulsion (Figure 1A). Briefly, the oil phase containing the cargo of interest was combined with an organic solution containing PCL and BKC which was then combined to the aqueous phase under rigorous stirring. BKC concentrations, oil core volumes, and rate of stirring were altered to determine the optimal NC formulation (Table 1A-C). BKC concentration had the largest standard variation of the mean (StdEM) on both NC size (StdEM ± 640nm) and polydispersity index (PDI) (sem ± 0.12), followed by changes in oil volumes (StdEM ± 16 for NC size, StdEM ± 0.03 for PDI) and then stirring speeds (StdEM ± 12 for NC size, StdEM ± 0.02 for PDI) (Table 1A-C). This is consistent with previous studies that showed similar changes in relative nanoparticle size with similar changes in stir speed and oil volume.^41–43^ Out of all combinations tested the synthesis parameters chosen for the rest of the study for the W/O emulsion consisted of the following: BKC (20wt/wt% PCL), 60μl oil, and 850rpm stirring speed which yielded the smallest average NC diameter of 271 ± 7nm with an average PDI of 0.06 ± 0.03, respectively (Table 1A). Spherical shape diameter and relative monodispersibility were confirmed by SEM and TEM (Figure 1B, C). Furthermore, successful encapsulation of BODIPY, a hydrophobic fluorescent cargo, was confirmed via confocal microscopy (Figure 1D).

**Table 1:** Evaluation of W/O NC size and polydispersity index (PDI) with changing synthesis parameters, including (a) BKC concentrations, (b) oil core volumes, and (c) rates of stirring (n=3). Values are color schemed as followed; green – lowest average values, yellow – second or third lowest values if more than four values for NC size and PDI, orange – second highest value for NC size and PDI, and red – highest values.

| BKC (wt/wt%PCL) | Oil (μl) | Stirrer Speed (RPM) | Average NC Diameter (nm) | Average PDI |
| --- | --- | --- | --- | --- |
| 2 | 60 | 850 | 2520 ± 2190 | 0.53 ± 0.38 |
| 10 | 60 | 850 | 394 ± 17 | 0.29 ± 0.03 |
| 20 | 60 | 850 | 271 ± 7 | 0.06 ± 0.03 |
| 100 | 60 | 850 | 279 ± 18 | 0.09 ± 7E-3 |

(b)
| BKC (wt/wt%PCL) | Oil (μl) | Stirrer Speed (RPM) | Average NC Diameter (nm) | Average PDI |
| --- | --- | --- | --- | --- |
| 20 | 50 | 850 | 321 ± 25 | 0.15 ± 0.06 |
| 20 | 70 | 850 | 366 ± 14 | 0.08 ± 0.04 |
| 20 | 80 | 850 | 305 ± 4 | 0.03 ± 7E-3 |
| 20 | 90 | 850 | 311 ± 10 | 0.07 ± 0.03 |

(c)
| BKC (wt/wt%PCL) | Oil (μl) | Stirrer Speed (RPM) | Average NC Diameter (nm) | Average PDI |
| --- | --- | --- | --- | --- |
| 20 | 60 | 550 | 336 ± 21 | 0.14 ± 0.01 |
| 20 | 60 | 650 | 289 ± 2 | 0.10 ± 0.05 |
| 20 | 60 | 750 | 290 ± 1 | 0.09 ± 7E-4 |
| 20 | 60 | 950 | 286 ± 11 | 0.11 ± 0.02 |
| 20 | 60 | 1050 | 292 ± 1 | 0.05 ± 0.02 |

**Figure 1:**
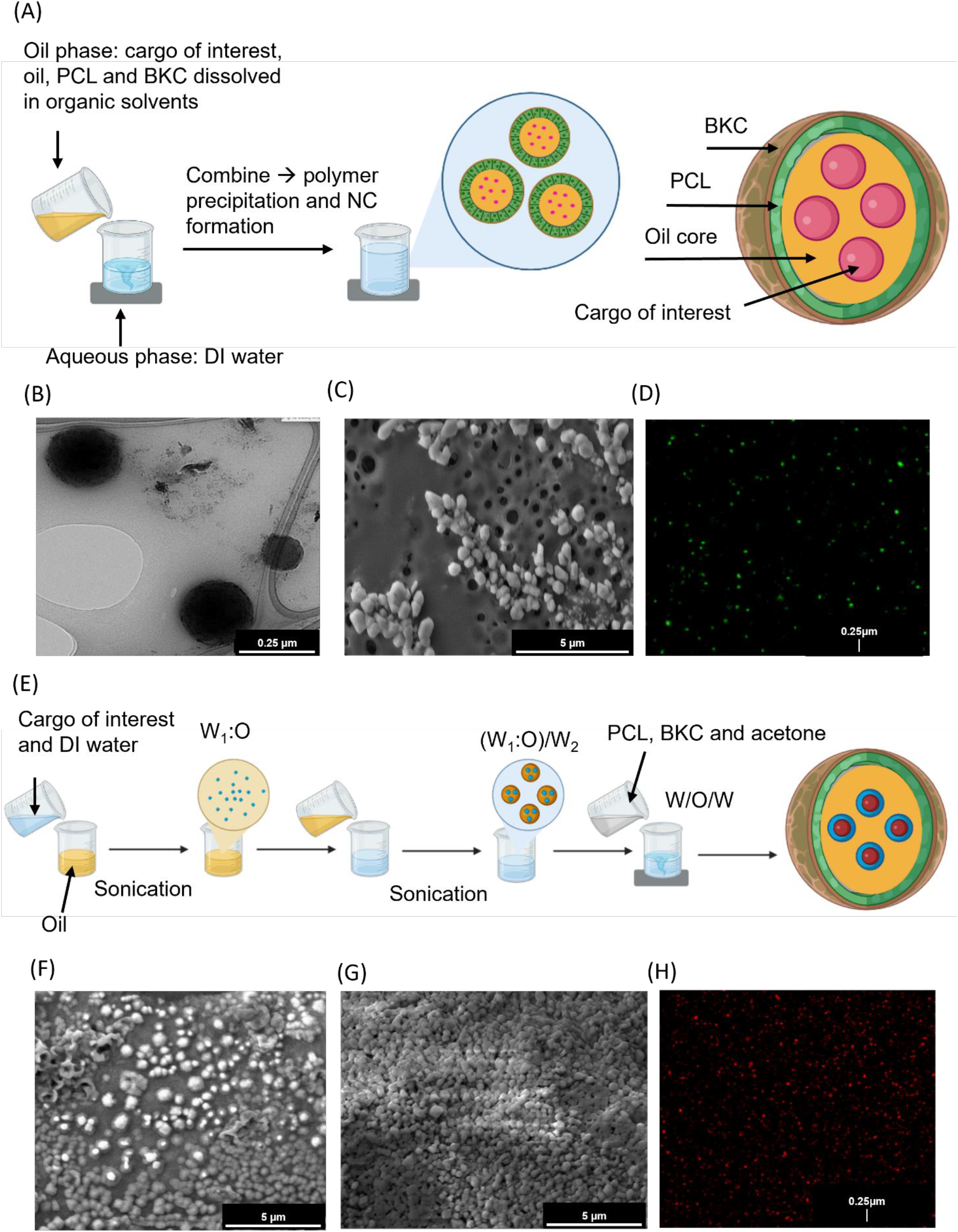
Water in Oil (W/O) and Water in Oil in Water (W/O/W) Nanocapsule (NC) synthesis. (A) Schematic outlining water in oil (W/O) synthesis of NCs where the cargo of interest is stored in an oil core surrounded by a polycaprolactone (PCL) polymer shell coated with cationic surfactant benzalkonium chloride (BKC), created in BioRender. The oil phase was combined with the aqueous phase under vigorous stirring for NC precipitation. Representative Transmission Electron Microscopy (TEM) (B) and Scanning Electron Microscopy (SEM) (C) images of blank (no cargo) W/O NCs. (D) Representative confocal image of W/O NCs loaded with BODIPY. (E) Schematic outlining water in oil in water (W/O/W) synthesis of NCs created in BioRender. The primary emulsion (W_1_:O) consists of the cargo of interest dissolved in DI water combined with the oil phase under sonication. The primary emulsion was combined with a secondary aqueous phase under sonication (W_1_:O)/W_2_. This was combined with an organic solvent containing the polymer and surfactant under rigorous stirring. Representative SEM images of (F) uncoated and (G) BSA coated blank W/O/W NCs. (H) Representative confocal image of W/O/W NCs loaded with Rh123. In images B-D and F-H the scale bar was added in post-processing to enlarge the text size using Image J.

For hydrophilic cargo encapsulation, NCs were synthesized via water in oil in water (W/O/W) emulsion (Figure 1E). For optimization of this fabrication method both W:O and (W_1_:O)/W_2_ ratios were altered for the W/O/W NCs (Table 2). These factors were varied to obtain the maximum NC yield while maintaining desired NC size. NCs formulated with a 25:75 W_1_:O and 10:90 (W_1_:O)/W_2_ had a significantly higher NC diameter from the other two treatments with a 10:90 (W_1_:O)/W_2_ phase (p=0.03 40:60 W_1_:O and p=0.02 50:50 W_1_:O, Table 2). This supports that varying the ratios of each phase can influence NC size. Like the W/O NCs, the average diameter and relative monodispersibility of coated and uncoated W/O/W NCs were confirmed by SEM (Figure 1F, G). Successful encapsulation of Rhodamine 123 (Rh123), a hydrophilic fluorescent cargo, for W/O/W NCs (Figure 1H) was also confirmed by confocal microscopy.

**Table 2:** Assessment of different W/O ratios and W/O:W ratios on W/O/W NC size and polydispersity index (PDI) (n=3). All ratios are reported as volumes in μl. Values are color schemed as followed; green – lowest average value, yellow – values within 10nm of lowest value for NC size and 0.01 for PDI, orange – values within 50nm of lowest value for NC size or 0.05 for PDI, and red – values over 50nm of lowest value for NC size or over 0.1 for PDI.

| W<sub>1</sub>:O Ratio | (W<sub>1</sub>/O):W<sub>2</sub> Ratio | Average NC Diameter (nm) | Average PDI |
| --- | --- | --- | --- |
| 25:75 | 5:95 | 363 ± 29 | 0.17 ± 0.02 |
| 25:75 | 10:90 | 309 ± 1 | 0.08 ± 0.02 |
| 40:60 | 10:90 | 275 ± 3 | 0.11 ± 0.02 |
| 40:60 | 25:75 | 282 ± 9 | 0.07 ± 0.03 |
| 50:50 | 5:95 | 338 ± 21 | 0.17 ± 0.01 |
| 50:50 | 10:90 | 280 ± 8 | 0.08 ± 0.02 |
| 50:50 | 25:75 | 287 ± 4 | 0.10 ± 0.02 |

Cargo were tested with varying charge, size, and hydrophobicity to assess if these parameters influenced NC stability, size, and encapsulation efficiency (EE). There was less than a 20nm increase in NC diameter for the W/O NC diameter when encapsulating either BODIPY or Cy5 while still maintaining a homogenous size distribution (Figure 2A, Table 3A) which is consistent with other studies.^44,45^ However, the Rh123 loaded W/O/W NCs had a 63nm diameter increase as well as slight increase in PDI (Table 3B). The bulky structure of Rh123^46^ could be attributing to the larger increase in NC size as another nanoparticle study showed about a 50nm increase with Rh123 NPs compared to the unloaded control.^47^ This data suggests NC diameter can be changed based on the size of the cargo being encapsulated. The EE of the BODIPY encapsulated W/O NCs was 68 ± 1.3% and 47 ± 0.7% for Rh123 encapsulated W/O/W NCs (Table 3C). The EE for BODIPY is similar to other NP fromulations^48^, however, Rh123 is much lower^49.^. The lower EE could be attributed to losing some of the solution during the transfer steps or the Rh123:PCL concentration was too high and there wasn’t enough PCL to encapsulate all the Rh123. Overall, this data supports our NCs can be designed to encapsulate either hydrophobic or hydrophilic cargo.

**Table 3:**
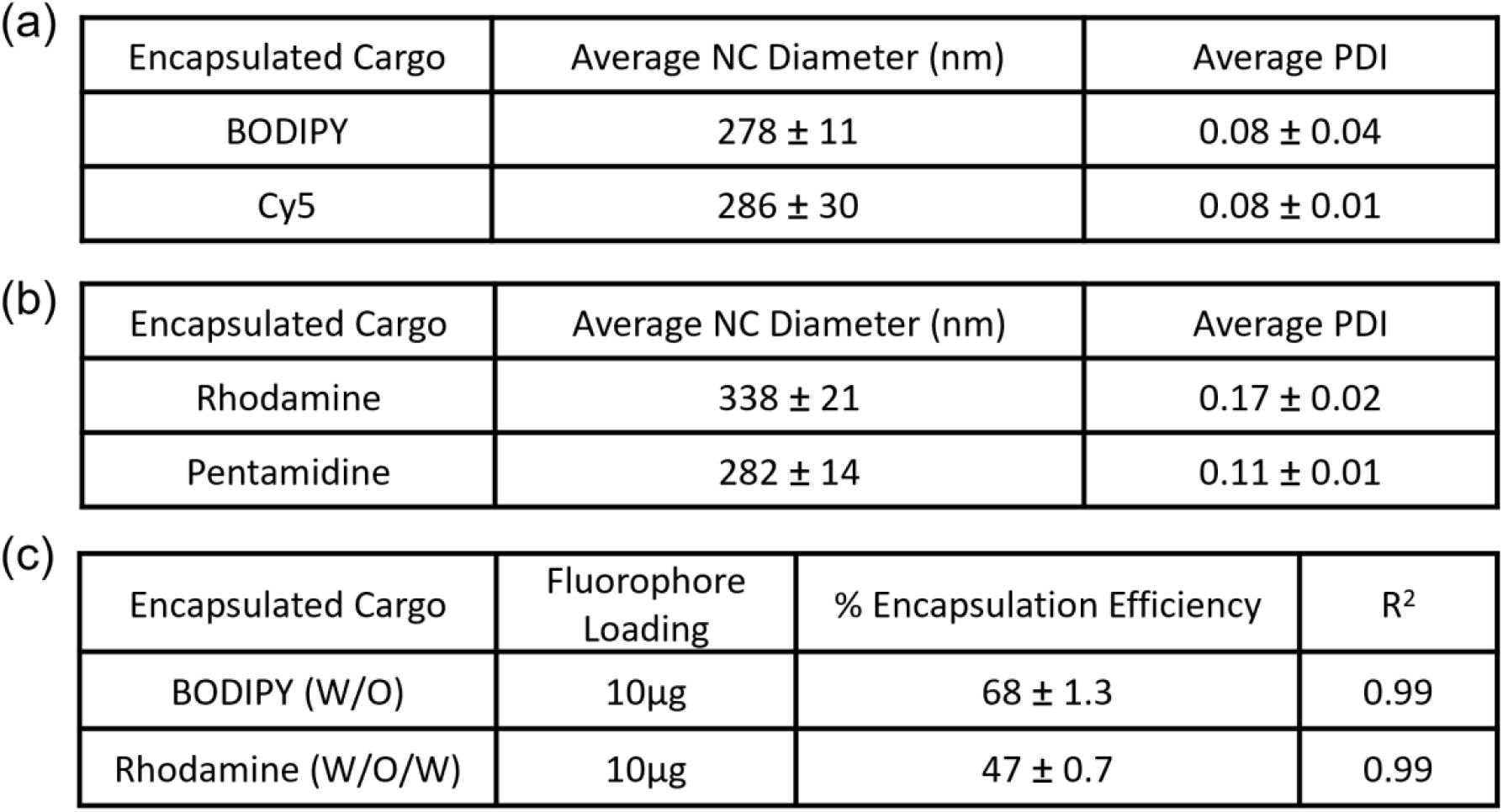
Influence of different encapsulated cargos on the size and polydispersity index (PDI) of (a) W/O NCS and (b) W/O/W NCs. (c) Percent of cargo that was successfully encapsulated (encapsulation efficiency) for BODIPY (hydrophobic) in W/O and rhodamine (hydrophilic) for W/O/W NCs (n=3-5). The coefficient of determination (R^2^) illustrates the fit of the standard curve to the actual encapsulation efficiency data.

|  |  |  |  |
| --- | --- | --- | --- |
| (a) | Encapsulated Cargo | Average NC Diameter (nm) | Average PDI |
| | BODIPY | 278 ± 11 | 0.08 ± 0.04 |
| | Cy5 | 286 ± 30 | 0.08 ± 0.01 |
| (b) | Encapsulated Cargo | Average NC Diameter (nm) | Average PDI |
| | Rhodamine | 338 ± 21 | 0.17 ± 0.02 |
| | Pentamidine | 282 ± 14 | 0.11 ± 0.01 |

|  |  |  |  |  |
| --- | --- | --- | --- | --- |
| (c) | Encapsulated Cargo | Fluorophore Loading | % Encapsulation Efficiency | $R^2$ |
| | BODIPY (W/O) | 10$\mu\text{g}$ | 68 ± 1.3 | 0.99 |
| | Rhodamine (W/O/W) | 10$\mu\text{g}$ | 47 ± 0.7 | 0.99 |

**Figure 2:**
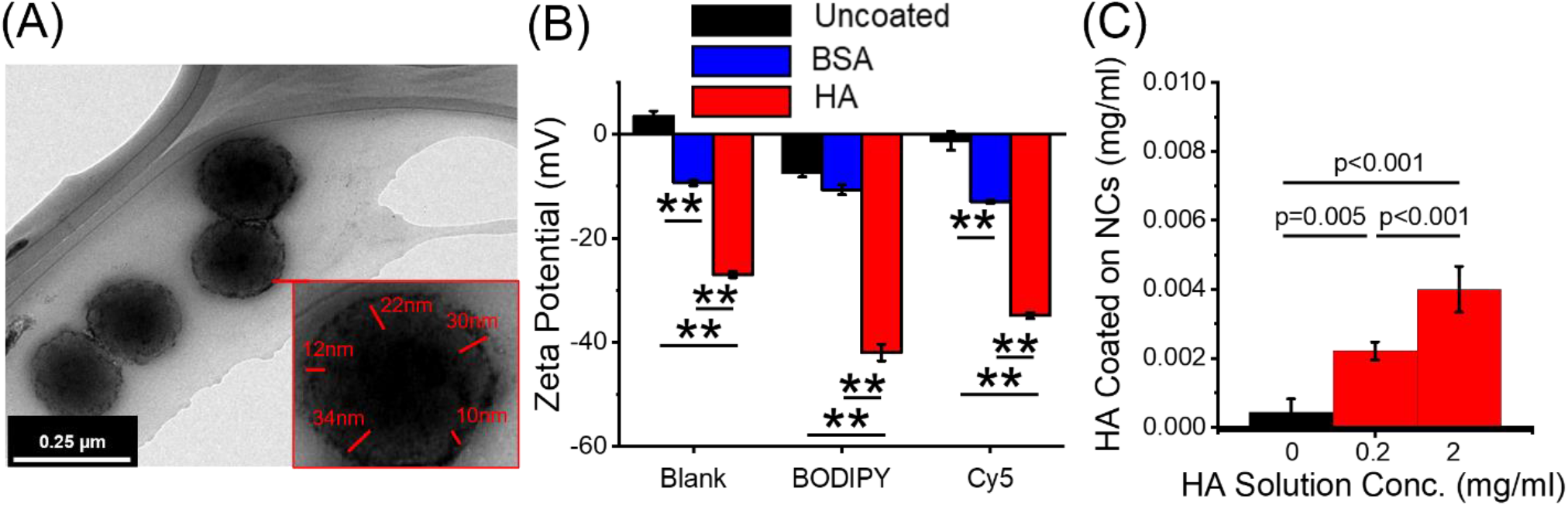
Various coatings can be applied to the outside of the NCs. (A) TEM image of BSA (5mg/ml) coated W/O NCs with a zoomed in NC highlighting the width of the BSA coating as measured in Image J. (B) Zeta potential of W/O NCs uncoated, BSA (5/mg/ml) and HA (0.2mg/ml) coated and either unloaded (blank) or loaded with BODIPY or Cy5 (n=3). (C) Concentration of HA in NC emulsions after coating with 0, 0.2, or 2mg/mL HA and washing for 10,000rpm for 10 minutes and re-suspending capsules in HA free buffer. **p<0.001, p,0.05 was considered significant as determined via ANOVA with Tukey’s post-hoc analysis.

### Anionic Coatings Can Adhere to NCs and Improve NC Stability

We next tested if BKC-PCL NCs could be coated with anionic coatings to improve NC stability and improve conjugations of different moieties for targeted delivery. Bovine serum albumin (BSA) and hyaluronic acid (HA) were chosen as they have different anionic strengths and can be used for *in vitro* or *in vivo* work. NCs were coated either with BSA or HA to determine if anionic compounds could adhere to the BKC/PCL framework.^36^ Compared to the average uncoated W/O NC diameter of 271 ± 7nm, HA coated NCs had an average diameter of 293 ± 4nm and BSA of 295 ± 1nm (Table 4). This equates to an average diameter increase of 22nm for HA and 24nm for BSA coated W/O NCs supporting anionic coatings do not greatly alter NC size. Scanning electron microscopy (SEM) and transmission electron microscopy (TEM) images both support the dynamic light scattering (DLS) measurements (Figure 1B, 1F, 1G, 2A). Furthermore, measurements were taken from the TEM images between the lighter shell (BSA coating) and darker core (NC) from the longest and shortest widths which averaged between 20-30nm supporting the increase in NC size displayed by DLS (Figure 2A, Table 4). All PDIs were under 0.1 and there was only a 0.01 difference between the coated and uncoated NCs demonstrating all formulations were monodispersed. The ability to adhere anionic coatings to the NCs allows the potential to attach various moieties to improve target specificity.

**Table 4:** Change in NC size and size distribution of uncoated and coated W/O synthesized NCs (n=3).

| Coating | Average NC Diameter (nm) | Average PDI |
| --- | --- | --- |
| None | $271 \pm 7$ | $0.06 \pm 0.03$ |
| HA (0.2mg/ml) | $293 \pm 4$ | $0.05 \pm 0.02$ |
| BSA (5mg/ml) | $295 \pm 1$ | $0.05 \pm 0.01$ |

W/O NC surface charge as measured by zeta potential was utilized to confirm if BSA and HA adhered to the outside of the NCs and assess stability of the coating. Furthermore, it was utilized to assess if encapsulated cargo, BODIPY (anionic) and Cy5 (cationic) significantly alters NC charge (Figure 2B). Although Cy5 loaded NCs did have a slight increase in zeta potential it did not significantly alter the NC charge compared to the blank W/O NCs (Figure 2B). This was consistent with a previous study where Cy5 itself was not a significant contributor to changes in zeta potential.^44^ Although the zeta potential of the uncoated BODIPY NCs was consistent with previous studies^45^ there was a significant negative charge compared to both the uncoated blank (p<0.001) and uncoated Cy5 W/O NCs (p=0.013, Figure 3B). When comparing the uncoated to both HA and BSA coated W/O NCs, all conditions tested except the blank BSA coated and BODIPY loaded BSA coated W/O NCs had a significant negative charge (p<0.001) ranging from −9 – - 12mV for BSA and −27 – −42mV for HA (Figure 2B). To further confirm that the NC coating was stably adhered to the surface of the NCs, we utilized a fluorescently tagged HA coated onto NCs at varying concentrations to confirm that HA remains adhered to the NCs after washing by centrifugation (Figure 2C). HA was detected on NCs after washing with centrifugation at 10,000rpm for 10 minutes and resuspending NCs in PBS. Additionally, increasing the HA solution concentration increased the amount of HA coated on the NCs. In addition, when comparing different BSA concentrations, the 10mg/ml BSA coated BODIPY encapsulated W/O NCs had a significant negative charge compared to other concentrations (p<0.001, S1B). These results are critical as charge can influence important factors such as cytotoxicity, cell permeability, and efficacy.^50–52^ The ability to tightly control surface charge allows the conjugation of ligands, peptides, antibodies, etc. for improved cell specificity making them a valuable drug delivery vehicle as well as screening tool for cell specific moieties for biological applications.

**Figure 3:**
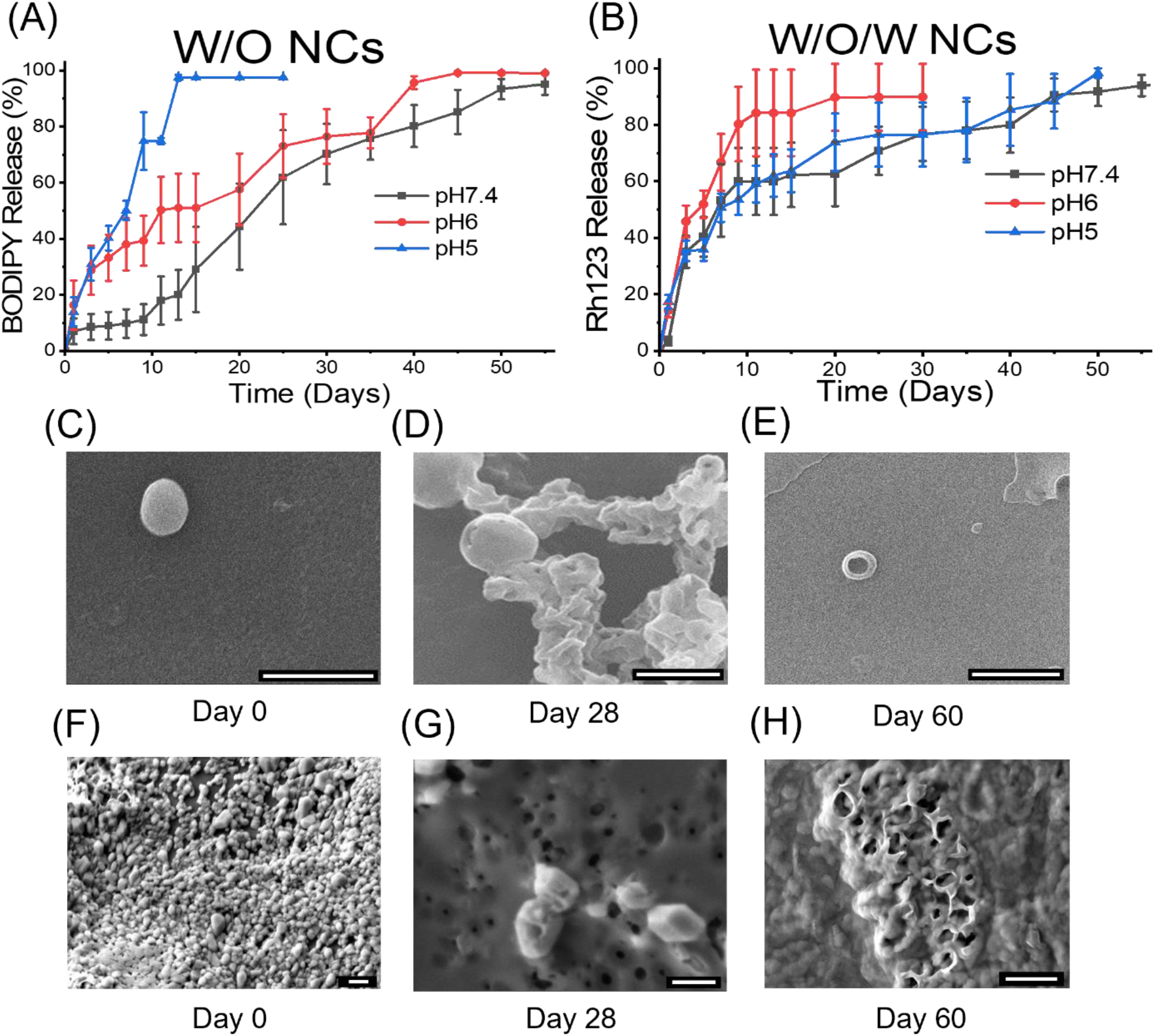
NCs can degrade and release cargos of interest at varying pHs. (A) *In vitro* release profiles of W/O NCs loaded with BODIPY and (B) *in vitro* release profiles of W/O/W NCs loaded with Rh123 in 1x PBS at pH 7.4, 6, and 5 (n=3-5). Release data are presented as the average value +/-standard error of the mean. All release data was normalized by subtracting the fluorescent value at time 0 and then dividing by maximum fluorescence value (100% release). Representative SEM images of W/O NC degradation (C-E) and W/O/W NC degradation (F-H) at day 0, 28 and 60 in IxPBS at pH 7.4. Scale bars are 1pm and were added in image post-processing using Image J.

### NCs Can be Freeze Dried and are Stable in Culture Media

Freeze drying is a method used to preserve NP integrity for extended periods of time and will allow translation of this technology into a clinical setting where storage for prolonged periods of time will be required.^53^ Therefore, we freeze dried NCs and assessed their stability over varying amounts of time. Mannitol was chosen as the cryoprotectant to help prevent aggregation and NC stability.^53^ Various mannitol concentrations were tested to determine the optimal concentration (S2A). There were no significant increases in NC size before and after freeze drying and reconstituting NCs in DI water between the different mannitol concentrations (S2A). Untreated NCs without cryoprotectant did not survive the freeze-drying process demonstrating a cryoprotectant is required for freeze drying (S2A). SEM images were taken before (S2B) and after (S2C) freeze drying to confirm DLS measurements. It is important to note the NC yield decreased after reconstitution which may be attributed to the mannitol crystallizing and imposing mechanical forces either breaking the NCs or forming aggregates.^53^ Future work will include investigating other cryo- or lyoprotectants as well as steric stabilizers which can improve NC stability.^54^

Lastly, the stability of uncoated, HA coated, and BSA coated W/O NCs were tested in Dubelco’s modified eagle medium (DMEM) culture media *in vitro* as well as being extracted and passed through a 29G 1cc insulin syringe. There was no significant difference in NC size or PDI after passing NCs through a 29G syringe as well as being incubated in DMEM for 1 and 24hr either at 37°C or room temperature (S3A-C). All PDIs for all conditions were under 0.12. Overall, the characterization data demonstrates the feasibility, stability and interchangeable capabilities for both charge and encapsulated cargo of the NCs. This supports the stability of NCs *in vitro* and delivery of NCs via IV injection through a 29G syringe for *in vivo* experiments.

### *In vitro* Cargo Release of W/O and W/O/W NCs

To determine the release profile of cargo from BKC-PCL NCs, BODIPY (W/O) and Rh123 (W/O/W) loaded NCs were placed in 1xPBS reservoirs at varying pHs and release rates were measured by fluorescence intensity of the solution. For the BOIPDY loaded W/O NCs the total average release time was between 45-50 days when incubating in PBS pH 7.4 (Figure 3A). There was no significant difference amongst any timepoint between pH 7.4 and pH 6. For pH 5 and pH 6 NCs, both had a similar increase in release rate in the first 24hr (Figure 3A). Although pH 6 and pH 5 NCs have a higher initial release rate compared to pH 7.4, there was only a significant difference in release between NCs incubating in ph 7.4 versus pH 5 on day six (p=0.004) and day seven (p=0.004) in the first week (Figure 3A). NCs in pH 5 almost have 100% of the cargo released by the end of the second week (Figure 3A). Furthermore, pH 5 had a significantly faster release profile compared to pH 7.4 every day during the second week (p=0.12, p<0.001, p=0.001, p=0.002, p=0.004, p<0.001, p=0.002) and only on the 13^th^ (p=0.04) and 14^th^ day (p=0.04) compared to pH6. There was no significant change for any timepoint in the release profiles for Rh123 loaded W/O/W NCs (Figure 3B); however, the release profile for W/O/W NCs was faster over the first week and then more gradual over progressive weeks compared to the W/O NCs. SEM images taken on day 0, 30, and 60 confirmed the degradation morphology of both the W/O and W/O/W NCs (Figure 3C-H). By day 30, pores begin to appear on the NC surface (Figure 3D, G) further supporting the release of fluorescent cargo as illustrated by the release studies (Figure 3A, B). By day 60, it appears all cargo has been released (Figure 3A, B) and only a partial PCL shell remains (Figure 3E, H).

These results support pH influences the degradation and in turn the release profiles of the W/O NCs. The faster degradation rate at lower pH is likely caused by increased hydrolytic degradation by random chain fission as this has been demonstrated as a major contributor in PCL degradation.^18^ Understanding this pH dependent mechanism is imperative because if the NCs are successfully endocytosed, they may be captured cellular organelles such as lysosomes or endosomes which have lower pH values (4-6) leading to faster drug release times.^55^

### NCs are Nontoxic and Can Selectively Target Insulin Producing sBCs and Human Islets *In vitro*

The pancreatic islet is a complex micro-organ comprised of multiple hormone secreting cells, the bulk of which are insulin producing β-cells. GLP-1R is expressed specifically on β-cells and the GLP-1R agonist Ex4 has been utilized extensively for quantifying and visualizing β-cell mass.^26,28^ ENTPD3 has recently been shown to be a marker for mature human and stem cell derived β-like cells; however, it has not been tested as a target for β-cell specific drug delivery.^56,57^ To assess the NCs ability to target β-cells *in vitro*, Cy5 encapsulated NCs were cultured with human islets from 2-3 independent donors for 24hr either uncoated or coated with BSA (5mg/ml), HA (0.2mg/ml), guinea pig IgG (1µl/ml), guinea pig ENTPD3 antibody (1µl/ml) or HA-Ex4 (0.2mg/ml). The islets were stained with the Zn^2+^ indictor FluoZin, as Zn^2+^ is specifically localized to insulin granules in β-cells, and NucBlue (live cell nuclei) and imaged to quantify Cy5 positive cells (Cy5+ or NC+) and β-cells as identified by Zn^2+^ positive staining (Figure 4A, S4A). Live cell imaging was utilized in lieu of fixed cells to quantify NC uptake as the fixation process would quench the fluorescence of the Cy5 in our NCs. After 24hr, both the ENTPD3 and HA-Ex4 coated treatments had significantly higher NC+ cells compared to the other control NC treatments (Figure 4B). Furthermore, at 48h after NC treatment there was an additional ∼6% and 10% increase of NC+ β-cells for HA-Ex4 and ENTPD3 coated NC, respectively, after 48hr incubation compared to 24hr (Figure 4B, S4B). To determine if the NCs were in fact targeting our cells of interest, in this case human β-cells, the NC+ cells were differentiated between Zn^2+^ positive (insulin+ β-cells) and Zn^2+^ negative (Figure 4C, S4C). The number of NC+ β-cells for the ENTPD3 and HAEx4 coated were significantly higher (P<0.001) than the other NC treatments at both 24 and 48hr (Figure 4C, S4C). There were no significant changes observed between all insulin-values for all treatments at both 24 and 48hr (Figure 4C, S4C). For the 48hr timepoint there was ∼11% increase in the NC+ β-cells for the ENTPD3 coated and ∼7% increase with the HA-Ex4 coated (Figure 4C, S4C). There were no significant changes observed between all insulin-values for all treatments (Figure 4C, S4C), suggesting that NCs coated with Exendin-4 or ENTPD3 antibody are highly human β-cell specific.

**Figure 4.**
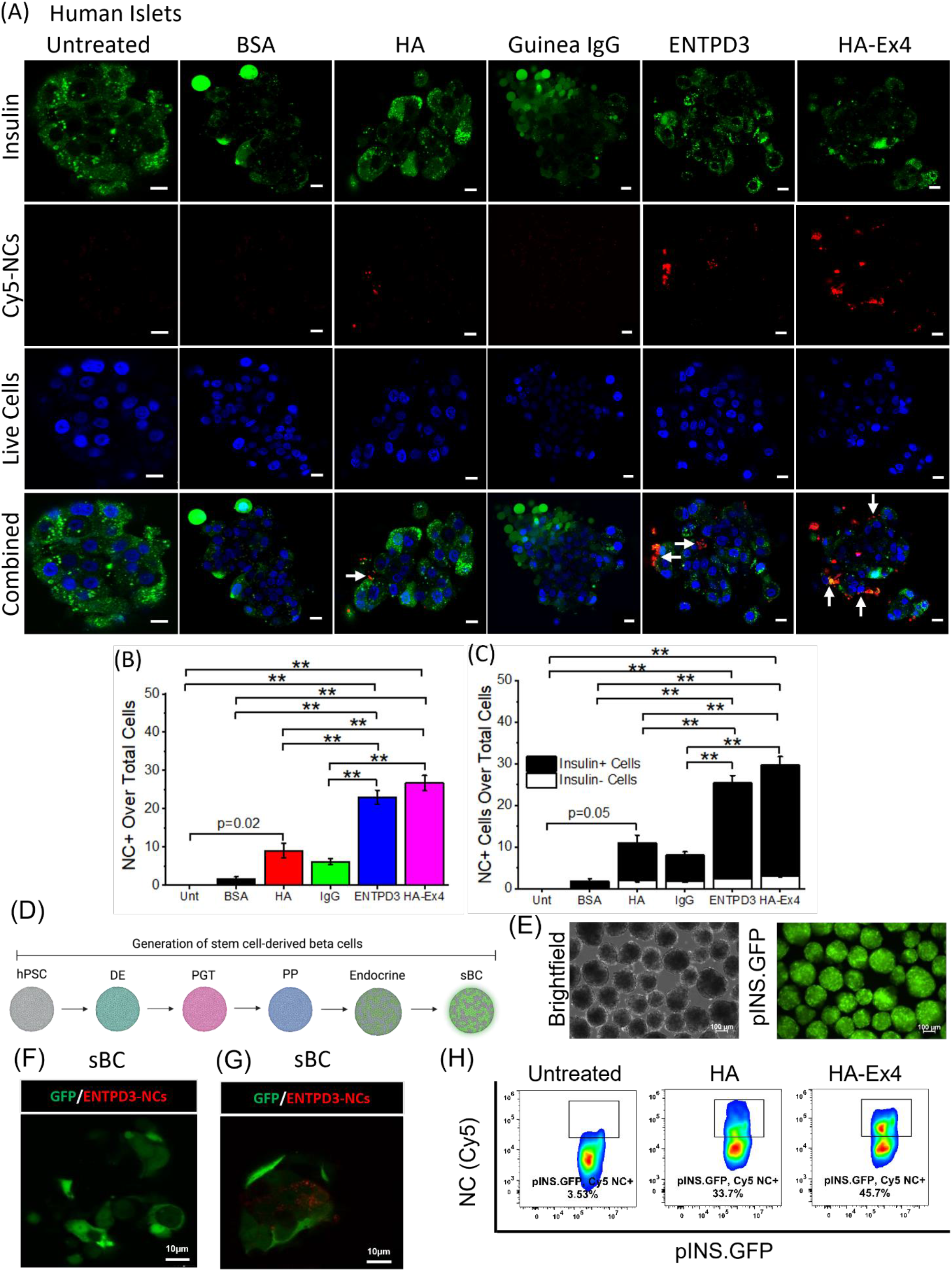
NCs specifically target insulin producing sBCs and human Islet cells in vitro. (A) Representative confocal images showing insulin (green), live cell nuclei (blue) and Cy5 NCs (red) with thevarious NC coatings. White arrows indicate NC+ and insulin+ cells. Scale bar is 10µm for all images. (B) Quantification of NC uptake with intact human islets. NCs were uncoated or coated with BSA (5mg/ml), HA (0.2mg/ml), guinea pig IgG (1µl/ml), ENTPD3 (lµl/ml), orHA-Ex4 (0.2mg/ml) and cultured with human islets for 24h (n=2-3). (C)Quantification of insulin NC+ and insulin+ NC+ cells with intact human islets for each NC treatment (n=2-3). **p<0.001, p<0.05 was considered significant as determined via ANOVA with Tukey’s post-hoc analysis. (D) Differentiation schematic to make sBC from human pluripotent stem cells. hPSC human pluripotent stem cell; DE – definitive endoderm; PGT – pancreatic gut tube; PP – pancreatic pro,enilor. (EJ representative live brightfield and plNS.GFP images of sBC on day 23 of differentiation protocol. Scale bar is 100 µm. Representative confocal image of (FJ untreated sBCs and (G) CyS loaded NCs with sBCs at 24hr (n=l). (HJ Flow cytometry quantification of CyS signal In live dispersed sBC co cultured with either HAor HA-Ex4 Cy5 loaded NCs at 24hr. Results were pre-gated on plNS.GFP+ cells(n=l).

Since human islets exhibit considerable variability between isolated preparations, we took advantage of a complementary source of functional human beta cells generated via direct differentiation from pluripotent stem cells. Stem cell derived beta cells (sBC) exhibit key β-cell characteristics, respond to glucose and rescue diabetes in preclinical animal models. We generated sBC from the embryonic human stem cell line Mel1^INS-GFP^ (Figure 4D). These cells have the benefit of a GFP reporter driven by the insulin promoter, allowing easy visualization of the insulin producing sBCs (Figure 4E). To determine if the NCs could specifically target insulin-producing sBCs, we co-cultured sBC clusters for 24 hours with uncoated or NCs coated with ENTPD3 antibody (1µl/ml) or HA-Ex4 (0.2mg/ml). Live cell imaging revealed Cy5+ sBC cells, identified by GFP expression, only in ENTPD3 coated NCs cultures further providing evidence for human β-cell specific targeting using NCs. (Figure 4F, G). To confirm these results, we also performed flow cytometry on sBC co-cultured with either HA or HA-Ex4 Cy5 NCs for 24h. Assessing our β-cell fraction using the pINS.GFP reporter, we observed that the addition of Ex4 on the surface of our NCs increased the percentage of sBC that took up Cy5 labeled NCs from 33.7% with HA-only coated NC controls to 45.7% with HA-Ex4 coated NCs (Figure 4H). Despite the significant levels of non-specific NC uptake with HA-coating, addition of Ex4 targeting improved uptake by 12% specifically in insulin expressing cells. Expression of the hyaluronan receptor on many cell types facilitates endocytosis of HA from the extracellular matrix.^58^ While it is unknown if sBC and human islets express this receptor, this may account for the significant non-specific uptake of HA-coated NCs compared to the IgG coated capsules in Figure 4B and C. Overall, our results strongly support that NCs coated with targeting peptide can be endocytosed by human β-cells with a high level of specificity.

### NCs Can Deliver Cargo of Interest to Insulin Producing sBCs and Human Islets *In vitro*

After demonstrating ENTPD3 antibody coated Cy5 containing NCs are taken up by sBCs, we then assessed if NCs can deliver additional cargo of interest to sBC clusters and human islets. To rapidly and effectively test β-cell specific targeting, we utilized the β-cell toxin pentamidine (PTM) as cargo. Empty (blank) or PTM loaded NCs were left uncoated or coated with BSA (5mg/ml), HA (0.2mg/ml), ENTPD3 antibody (1ul/ml) or HA-Ex4 (0.2mg/ml). Untreated and 1μM free PTM (no NCs) treated sBC clusters were utilized as additional negative and positive controls. sBCs and human islets were stained to observe live (NucBlue) and dead cells (PI) (Figure 5A,B). To identify β-cells, human islets were also stained with Fluozin3 to identify the zinc rich insulin granules in β-cells (Figure 5B), and in sBC we utilized the pINS.GFP reporter as previously described (Figure 5A). As expected, the positive control 1μM free PTM treatment killed the majority of cells in sBC clusters within 24hr (Figure 5C) while untreated samples exhibit low percentages of dead cells.^59^ Within 24hr of NC incubation with the sBC clusters, PTM loaded ENTPD3 coated NCs had a significantly higher percentage of dead cells compared to the untreated control (p=0.003, Figure 5A, C). A significant increase in cell death was observed with the PTM loaded ENTPD3 coated NCs compared to the blank BSA coated (p=0.008), blank ENTPD3 coated control (p<0.001), and BSA coated PTM loaded NCs (p=0.01, Figure 5C). Significant cell death was also observed at 72hr when comparing the PTM loaded ENTPD3 coated NCs to the untreated control (p=0.05), blank BSA coated (p=0.008), blank ENTPD3 coated (p=0.008), and BSA coated PTM loaded NCs (p=0.008, S5A, S5C, S5D). Except for the ENTPD3 coated PTM loaded treatment, there was no significant change in sBC viability between the untreated sBCs to all other NC treatments (Figure 5C, S5A, S5C, S5D), strongly supporting that the NCs are nontoxic and the amount of BKC on the NCs is below the cytotoxicity threshold. Further analysis was conducted where the number of GFP+ and GFP-dead cells were quantified as GFP correlates to insulin positive cells. The free PTM treatment could not be quantified for GFP+ and GFP-sBCs due to too much loss of cellular structural integrity. For the dead sBC clusters, 76% at 24hr (Figure 5D) and 69% at 72hr were GFP positive (S5D), supporting a high level of β-cell specificity and targeting in sBC. At 24hr there was a significant increase in GFP positive dead cells with PTM loaded ENTPD3 coated NCs compared to the untreated (p=0.008), blank BSA coated (p=0.004), blank ENTPD3 coated (p=0.01), and BSA coated PTM loaded NCs (p=0.008, Figure 5D). The increase in percentage of GFP positive dead cells was also significant at 72hr when comparing the PTM loaded ENTPD3 coated NCs to the untreated (p=0.05), blank BSA coated (p=0.008), blank ENTPD3 coated (p=0.03), and BSA coated PTM loaded NCs (p=0.008, S5D). Collectively, this data indicates that BKC-PCL NCs are non-toxic and can be designed to target and deliver cargo to the cells of interest to sBC with high specificity and efficacy over 24-72h.

**Figure 5:**
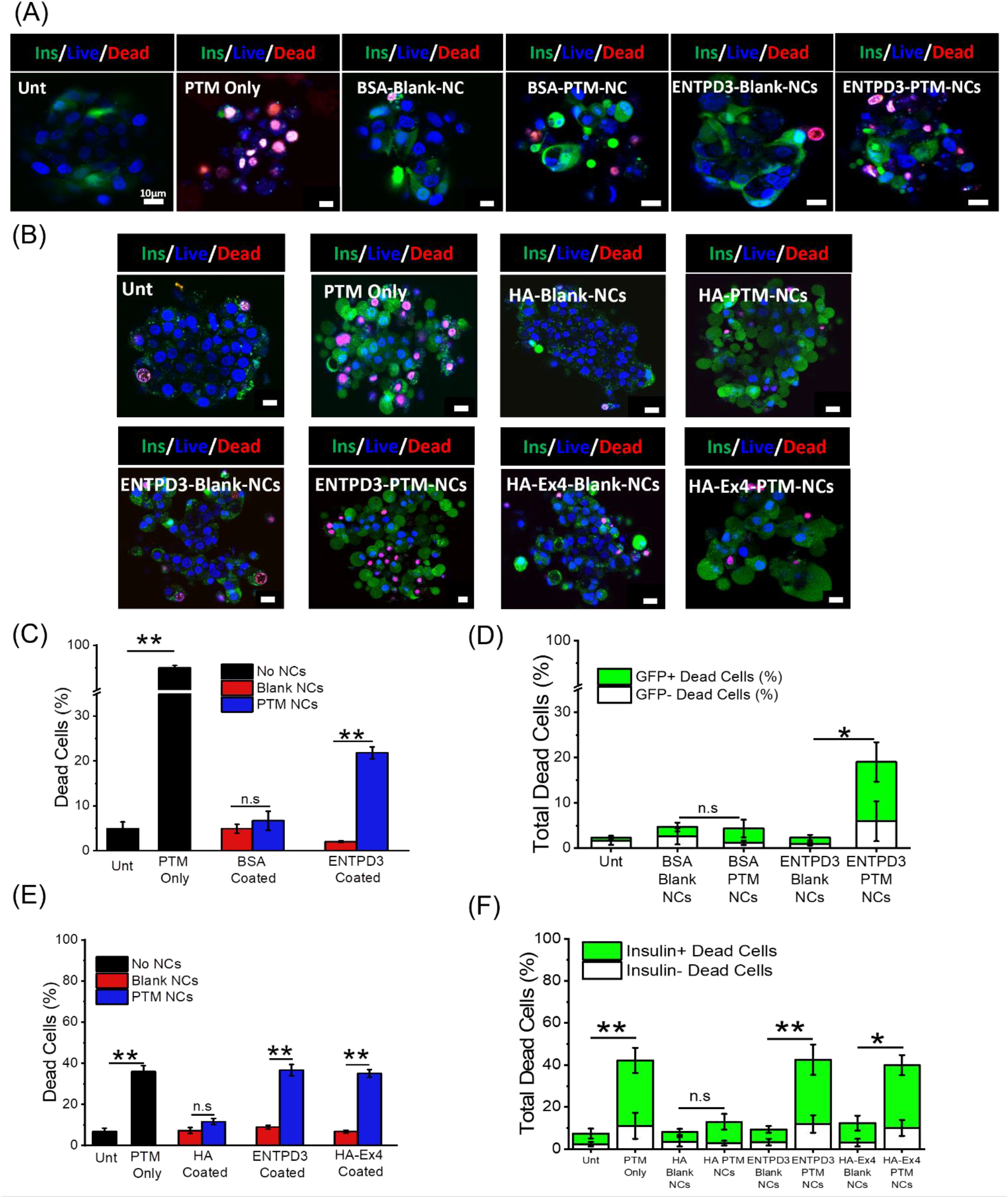
NCs can target, deliver, and release cargo of interest to insulin producing sBCs and human islets in vitro. (A) Representative confocal images of insulin+ (green), live (NuBlue, blue) and dead (Pl, red) sBCs either untreated, or treated with pentamidine (PTM) only, blank BSA coated NCs, PTM loaded BSA coated NCs, blank ENTPD3 antibody coated NCs or PTM loaded ENTPD3 coated NCs. (B) Representative confocal images of human islets either untreated or treated with PTM only, blank HA coated NCs, PTM loaded HA coated NCs, blank ENTPD3 antibody coated NCs, PTM loaded ENTPD3 coated NCs, blank HA-Ex4 coated NCs, or PTM loaded HA Ex4 coated NCs at 24hr. (C) Percentage of total dead sBCs for each NC treatment at 24hr and a breakdown of insulin+ versus insulindead cells (D) (n=4-6). (E) Percentage of total dead human islets for each NC treatment at 24hr and a breakdown of insulin+ versus insulindead cells (F) (n=3-4). p<0.05 via ANOVA with Tukey’s post-hoc analysis, **p<0.001, *p<0.01. P values displayed in (D) and (F) are for the insulin+ cells. No significance was observed between the insulin-treatments. Scale bar is l0µm and was added in post-processing using Image J.

The set of experiments described above was then repeated with human islets. Like the sBCs the PTM loaded ENTPD3 coated and HA-Ex4 coated had a significantly higher percentage of cell death compared to their blank controls at both 24 and 72hr (p<0.001, Figure 5E, S5E). The HA coated NCs had no significant change in the percentage of dead cells at both 24 and 72hr (Figure 5E, S5E). Furthermore, both the PTM loaded ENTPD3 and HA-Ex4 coated NCs had significantly more insulin+ dead cells compared to their blank controls at 24 (p<0.001, p<0.01) and 72hr (p<0.001) (Figure 5F, S5F). Of interest, NCs targeted with either Ex4 or ENTPD3 showed similar levels of NC uptake, PTM-induced cell death, and β-cell specificity indicating that ENTPD3 is a viable β-cell specific targeting peptide. Overall, these findings demonstrate these NCs can encapsulate, selectively deliver, and release a cargo of interest to both sBCs and human pancreatic β-cells with a high level of specificity.

### HA-Ex4 NCs Enrich in Pancreatic Islet β-cells *In vivo*

To test if Ex4 coated NCs could target pancreatic islets, Cy5 loaded HA coated NCS, or Cy5 loaded HA-Ex4 coated NCs were injected into the tail veins of 8 week old female NonObese Diabetic ShiLtSz-Prkdc^scid^ (NOD-scid) mice, an immunodeficient mouse model that does not develop T1D, and then sacrificed 24hr post injection. The staining of fixed pancreatic sections revealed an enrichment of NCs in pancreatic islets compared to the PBS control (Figure 6A). Overall, we found ∼ 9.4% of all β-cells positive for NCs as indicated by Cy5 co-localization with insulin staining in mice treated with HA-Ex4 coated NCs, while only ∼1.9% of β-cells were positive for NCs in mice treated with HA coated NC, a ∼5 fold enrichment in NC uptake(p=0.014, Figure 6B). Overall, the results of this study strongly support NC stability in the blood after tail vein injection and show proof of principle targeting to the mouse β-cell *in vivo*.

**Figure 6:**
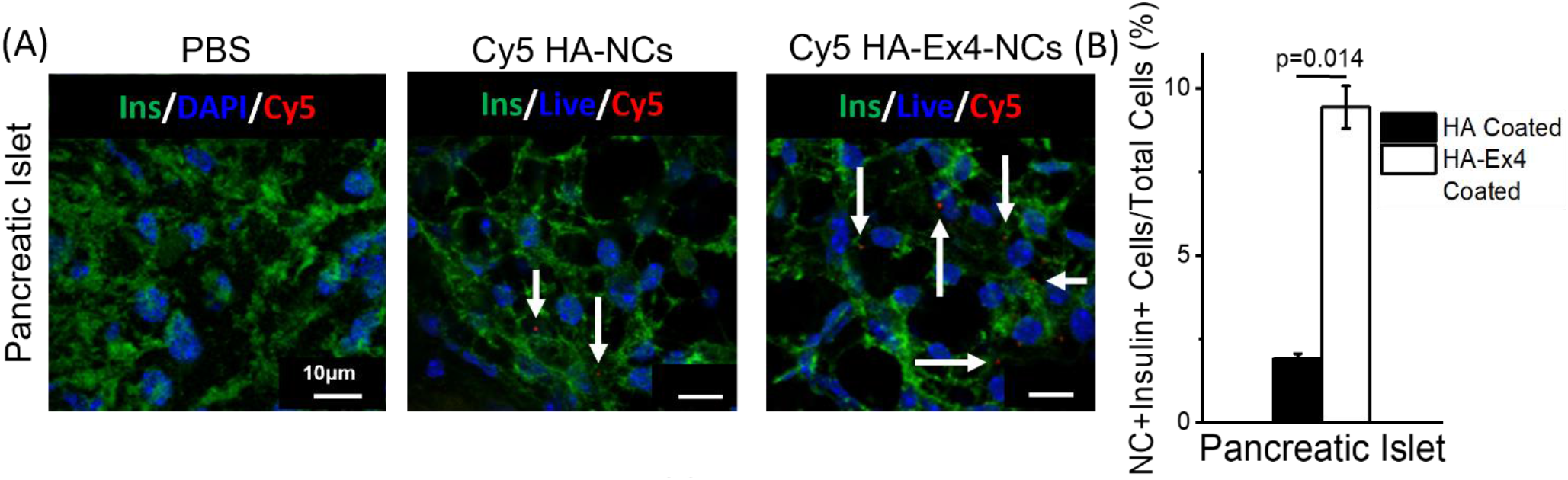
NCs can target pancreatic islet β-cells in vivo. (A) Representative confocal images of frozen pancreatic tissue sections from NOD-scid mice injected either with PBS only, Cy5 loaded HA-coated NCs, and Cy5 loaded HA-Ex4-loaded NCs. Tissue was stained for insulin (green), cell nuclei with DAPI (blue) and CyS loaded NCs (red). White arrows indicate CyS NCs. Scale bar is 10μm. (B) Quantification of percentage of NC+ cells per total cells in fixed pancreatic islet tissue (n=3). p<0.05 via ANOVA with Tukey’s post-hoc analysis. Scale bar is 10μm and was added in post-processing using Image J.

## CONCLUSION

In summary, in this study we report the development and characterization of a stable, non-toxic NC drug delivery vehicle targeted to the human β-cell. We demonstrated that NCs coated with 2 different targeting peptides are taken up by human β-cells with high specificity in 2 complimentary human islet models *in vitro*. Additionally, we demonstrated that NCs can deliver both hydrophilic and hydrophobic cargo to human β-cells. Finally, we have shown proof of principle stability and β-cell targeting of NCs injected into the tail vein of NOD-Scid mice *in vivo*. Our unique NC design allows for the interchangeable coating of targeting peptides for future screening of targets with improved cell specificity. The ability to target and deliver therapeutics to human pancreatic β-cells opens avenues for improved therapies and treatments to help the delay onset, prevent, or reverse T1D.

## MATERIALS AND METHODS

### Materials

PCL MW ∼14,000 (cat. #440752), BKC (cat. #12060), coconut oil (cat. #C1758), acetone (cat. #270725), bovine serum albumin (BSA) (cat. #A2153), pentamidine isethionate salt (P0547), divinyl sulfone (DVS, cat# V3700), Tris(2-carboxyethyl phosphine hydrochloride (TCEP, cat. #75259), and propidium iodide (cat. #P4170, Sigma) were all purchased from Sigma Aldrich (Milwaukee, WI). 4,4-difluoro-3a, 4a-di-aza-s-indacene fluorophore (BODIPY) (cat. #D3792), rhoda-mine-123 (Rh123) (cat. #cR302), NucBlue (cat. #R37605), 96 well plates (cat. #07-200721), Dulbecco’s Modified Eagle Medium (DMEM) (cat. #11-965-118), fetal bovine serum (FBS), penicillin, streptomycin, Dialysis tubing (21-152-8), PBS tablets (cat. #BP2944100), FluoZin™-3, AM, cell permeant (cat. # F24195), Guinea pig IgG isotype (IgG) (cat. # NBP1970365) and nanosphere size standards (cat. #09-980-027) were purchased from Fisher Scientific (Pittsburgh, PA). Lacey formvar/carbon-coated copper grid (cat. #01883-F, Ted Pella, Redding CA), hyaluronic acid (HA) (cat. #GLR001, R&D systems, Minneapolis, MN), ethanol (cat. #111000200, Pharmco). insulin syringes (cat. #CMD2626, BRANDZIG), cuvettes (part #759071D, Brandtech). Ectonucleoside triphosphate diphosphohydrolase-3 (ENTPD3) antibodies guinea pig anti-human NTPDase3 (hN3-2_c_) and mouse anti-human NTPDase3 (hN3-B3_s_) were purchased from Centre de Recherche du CHU de Québec – Université Laval CHUL (Québec, Canada). Dapi-Flu-oromount-G™ (cat. #17984-24) was purchased from Electron Microscopy Sciences (Hatfield, PA). Insulin primary antibody (cat. #ab63820) was purchased from Abcam (Cambridge, MA). Anti-rabbit secondary antibody Alexa Fluor® 488 anti-IgG donkey (cat. #102649-730) was purchased from VWR (Radnor, PA).

### Nanocapsule Synthesis

PCL-NCs were synthesized via a water in oil (W/O) emulsion technique for encapsulating hydrophobic cargo using methods adapted from previously described protocols (Figure 1A).^15,52^ The oil phase consists of two solutions. The first is coconut oil (60μl) dissolved in ethanol (750µl) and the second is PCL (2mg/ml) and BKC (20wt/wt% PCL) dissolved in acetone (4.25ml). Both solutions were combined and stirred briefly and then immediately poured into the aqueous phase, consisting of 10mL deionized water for 1-2min with rigorous stirring. The ethanol and acetone were removed using a rotavapor (R-100, Buchi, New Castle, DE), leaving a NC suspension in DI water. NCs were washed twice by centrifugation at 20,000rcf for 10min and resuspending the NCs in fresh DI water. Various BKC concentrations, oil volumes, and stirring speeds were tested to identify the optimal synthesis conditions for the generation of stable NCs, as outlined in Table 1. W/O synthesized BKC-PCL-NCs were loaded with 10ul of either BODIPY or Cy5 dissolved in the ethanol solution and the volume of cargo added was subtracted from the amount of ethanol added allowing the total volume to remain at 750ul. Coated W/O NCs incubated either with BSA of HA for at least 30min at room temperature and then washed via centrifugation as previously stated above. All parameters were tested using at least three separate NC batches.

A water in oil in water (W/O/W) double emulsion-solvent evaporation synthesis was used to encapsulate hydrophilic cargo in NCs similar to a protocol previously described (Figure 2A).^60^ The primary emulsion (W_1_:O) phase consisted of hydrophilic cargo dissolved in DI water (W_1_) and coconut oil (O) at a ratio of 40:60. The DI water with the dissolved cargo was combined with the coconut oil and sonicated at 20% power for 3 min on ice. The W_1_:O emulsion was then combined with DI water (W) at a 1:10 ratio and sonicated at 20% power for 3min (W/O/W). A solution containing PCL (2mg/ml) and BKC (20wt/wt% PCL) dissolved in 4.25ml acetone was added dropwise to the W/O/W emulsion under vigorous stirring for 2min. The acetone was evaporated and the NCs were washed as described above. The W_1_:O phase was tested at various ratios ranging from 25:75 – 50:50 and ratios of the (W_1_:O)/W_2_ emulsion were tested at volume ratios from 5:95 – 25:75 (Table 2). Coating the W/O/W NCs either with BSA or HA was the same as stated above. All parameters were tested in triplicate as the (W_1_:O) NCs stated above.

### Nanocapsule Characterization

#### NC size, polydispersity, and zeta potential

The mean particle diameter and PDI of the NCs were measured via dynamic light scattering (DLS) and the overall charge was measured via zeta potential. Both DLS and zeta potential were measured either with a ZetaPALS (Brookhaven Instruments, Holtsville, NY) or a Malvern Zetasizer (Malvern Panalytical Ltd, UK). DLS measurements were taken of uncoated, BSA (5mg/ml) coated, and HA (0.2mg/ml) coated W/O NCs. Zeta potential measurements of W/O NCs were taken of blank, BODIPY loaded, and Cy5 loaded either uncoated and coated at various concentrations ranging from 2mg/ml – 25mg/ml BSA and 0.2mg/ml – 5mg/ml HA (Figure 3B, S1A, B). Samples were diluted with DI water until the solution was clear and 3ml were placed in polystyrene spectrophotometry cuvettes for analysis. Each experimental condition was performed with three separate W/O batches.

The mean particle diameter and PDI of blank, Rh123 (0.06mg/ml) loaded, and pentamidine (PTM, 50mg/ml) loaded W/O/W NCs were also measured via DLS as stated above. Each experimental condition was performed with three separate W/O/W batches.

#### Electron Microscopy

DLS size measurements were confirmed via transmission electron microscopy (TEM) using a FEI Technai T12 transmission electron microscope (ThermoFisher Scientific, Waltham, MA) and scanning electron microscopy (SEM) using either a JEOL JSM 7000F field electron scanning electron microscope (Akishima, Japan) or a TESCAN S8252G RAMAN SEM/FIB (Brno, Czech Republic).

For TEM, 2μl of NC suspension was mounted on a lacey formvar/carbon-coated copper grid and air dried before imaging. For SEM, samples were air dried on carbon tape and were gold sputtered before imaging and imaged between 5 – 15kV.

#### Confirmation of NC coating

NCs were coated with 5mg/ml BSA or 0.2mg/ml HA as described in the NC Synthesis. Their size, PDI, and zeta potential were then measured using the above DLS instruments and techniques. Additionally, TEM assessed NC size and provided visual evidence supporting NC coating. To further quantify the amount of HA coated onto the NCs and assess the stability of NC coating after centrifugation. NC were prepared as described above, then diluted down to an optical density (OD) of 20 ± 0.48 measured at a wavelength of 250 nm.

NCs were incubated with 0.2 mg/mL or 2 mg/mL TMR-HA for 30 minutes with stirring at 200 rpm at room temperature, after which 1 mL aliquots were centrifuged at 10,000 rpm for 10 minutes. The supernatant was decanted, and the remaining NCs were diluted with DI water up to 1 mL. This washing step was repeated 2 times to remove the excess TMR-HA that was not coated on the NCs. A sixteen-point TMR-HA standard curve was generated which ranged in concentration from 0 mg/mL – 0.5 mg/mL. Samples and standards, 200 µL each, were added to a black-walled, clear-bottom 96-well microplate in triplicate. The microplate was read with a Synergy H1 microplate reader (BioTek) with an excitation wavelength of 546 nm and an emission wavelength of 576 nm with a Gain of 50. The fluorescent intensity of the standard curve was fit to the following equation in MATLAB, y=ae^bx+ce^dx. The coefficients generated from the line of best fit were used to solve for the concentration of the TMR-HA coated onto the NCs. (n=3)

#### Encapsulation efficiency (EE) of NCs

BODIPY (W/O) or Rh123 (W/O/W) were encapsulated in BKC-NCs as described above. After synthesis and washing the NCs 2-3 times with DI water, the NC suspensions were dissolved in acetone and 250μl samples were placed in triplicate in a 96 well plate. Standard curves were created by dissolving all components utilized for NC synthesis (excluding the DI water, acetone, and ethanol) in the same volume of acetone to emulate 100% encapsulation efficiency (EE) and diluted by factors of 1:2 – 1:1,000 to generate a curve with a linear correlation between concentration and fluorescence. From the linear equation generated, all standard curves had a coefficient of determination (R^2^) of 0.99 supporting the interpolation of samples. The following equation was then used to calculate the EE:

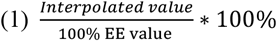

Fluorescent intensities of the samples were measured at 485/515nm for BODIPY and 500/530nm for Rh123 using a Synergy H1 microplate reader (Agilent Technologies, Santa Clara, CA) and analyzed via Gen5 3.11.19 software. Samples and standards were measured without a lid and set at a read height of 7mm with a 100msec delay, xenon flash, light source, and a gain of 100. All samples were read on the same plate at the same time as the standard curve samples to minimize error between different plate readings.

#### Freeze Drying NCs

Blank NCs at an OD 20 were suspended in 5-10ml of either DI water or 2.5w/v, 5w/v, or 10w/v mannitol solutions. All solutions were stored in −80C overnight and then lyophilized for 24 – 48hr. Freeze dried NCs were stored at room temperature protected from light for one month. NCs were then resuspended in DI water and centrifuged at 20,000rcf for 10min. The mannitol solution was discarded and replaced with fresh DI water. NC size was analyzed by DLS and SEM (S2A-C).

#### Assessment of NC Stability in vitro

The stability of both coated and uncoated NCs was tested both in culture media and after being passed through a 29G 1cc insulin syringe. Uncoated, BSA (5mg/ml) and HA (0.2mg/ml) W/O NCs were incubated in DMEM culture media with 10% FBS and 5% penicillin/streptomycin for 1hr and 24hr both room temperature and in an incubator set at 37°C and 5% CO_2_. For the syringe transfer, 300μl of uncoated, BSA (5mg/ml) and HA (0.2mg/ml) coated W/O NCs were taken up and passed through a 30G insulin syringe needle. NC size and PDI were measured via DLS before and immediately after each treatment. Three separate W/O NC batches were tested (S3A-C).

### *In vitro* Cargo Release

BSA coated NCs loaded with either BODIPY or Rh123 were placed in a dialysis bag (12,000 – 14,000 MWCO) in 415ml of 1x phosphate buffer saline (PBS). The beaker was sealed with parafilm to mitigate evaporation. The beaker was placed on a stir plate at 100 rpm, protected from light and stored at 37°C. 1ml samples were withdrawn every 24hr for the first 15 days then once every 5 days until day 60. Experiments were performed at pH of 7.4, 6, and 5, where the pH of each reservoir was checked weekly and adjusted with either 1M HCL or NaOH as necessary. Fluorescence intensity of samples was measured at 485/515nm for BODIPY and 500/530nm for Rh123. Samples were measured in triplicate. The intensity at each timepoint was normalized by the 0-time point fluorescence intensity and the normalized value was averaged across experimental replicates. Percent release of the fluorophore was calculated by the following equations:

Difference between fluorescence intensity at time 0 from each timepoint *I*_*N*_(*t*):

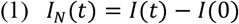

Percent release at each timepoint:

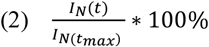

Where, I(t) is the fluorescent intensity at a given time (t), I(0) is the intensity at time 0, and t_max_ is the time where the fluorescence intensity was at the maximal value. SEM imaging was utilized to visualize the degradation of NCs at day 0, day 28, and day 60.

### Selectivity and Cytotoxicity of β-cell Targeted NCs *In vitro*

#### SBC differentiation

Human pluripotent stem cells (hPSCs) containing a green fluorescence reporter were cultured and direct differentiated as previously described.^61–63^ Briefly, hPSC cultures were maintained on Geltrex-coated plates in mTeSR+ media (STEMCELL Technologies). To induce direct differentiation hPSC cultures were transitioned into suspension-based bioreactor cultures for 72 hours. Differentiating suspension cultures were monitored daily. sBCs emerged approximately at d13 of the differentiation process and were utilized for experiments in this study between day 22-32.

#### Human cadaveric islets

Human islets from cadaveric donors were obtained through the Integrated Islet Distribution Program (IIDP). Donor islet viability, purity, and donor characteristics including age, sex, BMI, and race are shown in Table 5.

**Table 5:** Human cadaveric islet viability, purity, and donor characteristics for all donors received from IIDP.

| Donor ID | Viability (%) | Purity (%) | Age | Sex | BMI | Race |
| --- | --- | --- | --- | --- | --- | --- |
| SAMN34075901 | 91 | 83 | 39 | M | 41.8 | White |
| SAMN34130383 | 99 | 95 | 42 | F | 29.2 | Hispanic/Latino |
| SAMN34411471 | 94 | 92 | 45 | M | 33.1 | White |
| SAMN36704819 | 80 | 85 | 52 | M | 25.9 | White |
| SAMN36705973 | 96 | 95 | 36 | F | 30.2 | Black/African American |
| SAMN36823227 | 94 | 77 | 41 | F | 38.4 | White |
| SAMN37638596 | 99 | 80 | 34 | M | 27.2 | Native Hawaiian/Pacific Islander |
| SAMN38117428 | 95 | 95 | 47 | M | 28.1 | White |

#### Synthesis of Ex4 conjugated to HA (Ha-Ex4)

The modification of HA with DVS was prepared as previously described with some modifications.^64^ The low molecular weight HA was dissolved in 0.1 M NaOH at 2% w/v (∼200 μmol hydroxyl groups per mL). DVS was added into the HA solution on a magnetic stirrer at 1000 rpm at a molar ratio of 10x the hydroxyl groups of HA. The reaction was carried out for 10 min at room temperature and stopped by adjusting the pH to 5 using 1 M HCl. The solution was dialyzed (MWCO: 3.5 kDa) against deionized (DI) water (pH ∼5.3) every day with a new medium for 5 days and then freeze-dried. Cysteine modified Ex4 was provided by Dr. David Hodson and Dr. Johannes Broichhagen and was synthesized as previously described.^65^ Before conjugating the modified HA to Ex4, the Ex4-Cys was also modified with a reducing agent, TCEP at a molar ratio of 1:10 using DI at room temperature with stirring at 1000 rpm for 30 min and then freeze-dried.^66^ The modified dried forms of Ex4 and HA were re-dissolved in DI and the pH of the solution was adjusted to 9 with 1 M HCl. The solution was incubated at 37 °C, 1000 rpm for 12 hours in a nitrogen-purged flask on an oil-bath. To block the unreacted acryloyl groups in HA-Ex4 conjugates, 2 M excess of cysteine was added to the reaction solution. The pH of the reaction solution was adjusted to 9 again and incubated at 37 °C for another 12 hours. Then, the reaction was stopped by reducing the pH to 7.0 with 1 N HCl. HA-Ex4 conjugates were purified by dialysis (MWCO: 3.5 kDa) against DI water (pH ∼7.2) every day with a new medium for 5 days and then freeze-dried. *NC trafficking*. GFP-expressing sBC were cultured with Cy5 loaded NCs either uncoated or coated with a ENTPD3 antibody (1µl/ml) for 24hr in mTeSR+ media. Human islets were cultured with Cy5 loaded NCs uncoated or coated with BSA (5mg/ml), HA (0.2mg/ml), IgG (1µl/ml), ENTPD3 (1µl/ml), or HA-Ex4 (0.2mg/ml) for 24 and 48hr. Human islets were stained with FluoZin-3, a zinc sensor that localizes to insulin granules, and NucBlue a live cell dye. Clusters and human islets were imaged on a Leica STELLARIS 5 LIAchroic with a 40X water immersion objective, 405nm, 488nm, and 638nm solid state lasers and 3 HyD spectral detectors. NC positive (NC+) cells were quantified via manual counting in ImageJ, where any Cy5 signal surrounding individual nuclei stained by NucBlue were considered positive (NIH). NC negative (NC-) cells were defined as cells that did not have NCs on the periphery or within the insulin or NucBlue stain. Insulin positive cells were counted manually as determined by the presence of high fluorescence intensity staining of insulin granules, diffuse low intensity FluoZin-3 staining in the cytoplasm was not considered insulin positive.

#### Targeted delivery

Blank and PTM loaded NCs were coated either with an ENTPD3 antibody (1μl/ml) or BSA (5mg/ml) and added to sBC in culture medium for up to 168hr. SBC clusters were also treated with PTM only (1μm) as a positive control. Clusters were stained with NucBlue (live) and propidium iodide (dead, 10μl/ml). Clusters were imaged on a Leica STELLARIS 5 LIAchroic with a 40X water immersion objective, 405nm, 488nm, and 514nm solid state lasers and 3 HyD spectral detectors. Live/dead cells were quantified via manual counting in ImageJ (NIH). 4-5 different sBC differentiations were used and 3-6 sBC clusters were imaged per treatment per experiment.

#### Flow Cytometry

After co-culture with Cy5 NCs, wells with dispersed sBC were washed twice with 1xPBS to remove any NC not taken up by the cells. Cells were removed from the plate using TryPLE solution (Gibco 12604021) for 3 minutes and quenched with equal volume of media. Cells were filtered through cell strainer into FACS tubes, washed once with 1xPBS, and resuspended in 200 μL of FACS buffer before analysis on the Cytek Aurora 5L spectral cytometer. sBC cultured alone were used as compensation control for pINS.GFP expression and pure Cy5 NC were used as compensation control for NC signal. 40,000 events were collected for each sample.

### NC Trafficking *In vivo*

All animal studies were conducted at the University of Colorado Anschutz Medical Campus and were conducted in accordance with the guidelines and relevant laws set by the University of Colorado and the National Institutes for Health guide for the care and use of laboratory animals. All procedures were approved by the University of Colorado Institutional Animal Care and Use Committee under protocol #00929. Either PBS, Cy5 loaded HA coated NCs, or Cy5 loaded HA-Ex4 coated NCs were injected into the tail veins of 8-week old female non-obese diabetic/severe combined immunodeficiency (NOD-scid) mice. These mice were sacrificed 24hr post injection and the pancreata were extracted. The pancreata were cryosectioned in 10μm thick sections and placed on slides and immunohistochemistry was performed. Frozen sections were rehydrated, air dried, and then permeabilized with PBS containing 5% NDS and 0.25% Triton-X-100 for 5min. A blocking buffer consisting of 5% NDS and 0.1% Triton-X-100 was then applied to the sections for 5min. Sections were washed with PBS and then a PBS solution consisting of 5% NDS, 0.1% Trition-X-100 and an anti-insulin primary antibody (1:1,000) was applied and the tissues incubated either in 4C overnight or at room temperature for 2hr. Sections were rinsed three times with PBS and then a PBS solution containing 5% NDS and an anti-rabbit secondary antibody (1:500) was added to the sections and incubated at room temperature for 2hr protected from light. DAPI fluoromount (cell nuclei stain) was added to the sections and the slides were covered with a glass cover slip and sealed using nail polish. Pancreatic tissues were imaged on a Leica STELLARIS 5 LIAchroic with either a 40X water or 63x oil immersion objective, 405nm, 488nm, and 638nm solid state lasers and 3 HyD spectral detectors. NC positive and NC negative dead cells were quantified via manual counting in ImageJ (NIH). 8-10 images were taken per pancreatic tissue per treatment. 2-3 mice were used per treatment.

### Statistical Analysis

Statistical analyses were performed using Origin software (OriginLabs, Northampton, MA). Two sample t-tests and one-way analysis of variance (ANOVA) with Tukey’s post hoc analysis were performed as indicated. A p-value of <0.05 was considered statistically significant.

## ASSOCIATED CONTENT

### Supporting Information

Figures S1-S5.

## AUTHOR INFORMATION

## Author Contributions

N.L.F. and H.A.R. conceptualized and received funding for the project, oversaw experiments and data analysis. J.C., J.M.B., K.H., A.O., A.F., J.A., A.K., J.K.H., and A.S. performed experiments. J.C., J.M.B., K.H., A.O., A.F., J.A., and J.K.H. analyzed data. D.J.H. and J.B. provided experimental peptide and oversaw experimental design. J.C., J.M.B., A.F., J.A., D.J.H, J.B., H.A.R., and N.L.F. wrote and edited the manuscript. All authors have given approval to the final version of the manuscript.

## ACKNOWLEDGMENT

Funding for this work was provided by the following grants: NIH Diabetes Research Center Pilot and Feasibility Award to N.L.F. and H.R. through the NIDDK award P30-DK116073, JDRF 1-INO-2022-1231-S-B and JDRF 3-SRA-2023-1367-S-B to N.L.F. and H.R, and ADA 7-20-JDF-020 to N.L.F. Work in the lab of HAR is supported by NIH/NIDDK R01DK132387 and the JDRF 2-SRA-2023-1313-SB. The authors would like to acknowledge support from the Diabetes Research Center at the University of Colorado Anschutz Campus P30-DK116073 and the associated core facilities that were utilized in support of this work. D.J.H. was supported by MRC (MR/S025618/1), Diabetes UK (17/0005681 and 22/0006389) and UKRI ERC Frontier Research Guarantee (EP/X026833/1) Grants. This work was supported on behalf of the “Steve Morgan Foundation Type 1 Diabetes Grand Challenge” by Diabetes UK and SMF (grant number 23/0006627 to D.J.H. and J.B.). This project has received funding from the European Research Council (ERC) under the European Union’s Horizon 2020 research and innovation programme (Starting Grant 715884 to D.J.H.). This project has received funding from the European Union’s Horizon Europe Framework Programme (deuterON, grant agreement no. 101042046 to JB). We would also l like to acknowledge organ donors and their families that made this work with human islet from the IIDP possible.

## ABBREVIATIONS

ANOVA: analysis of variance
BKC: benzalkonium chloride
BSA: bovine serum albumin
DI: deionized
DLS: dynamic light scattering
DMEM: Dubelco’s modified eagle medium
DYRK1A: dual specificity tyrosine-phosphorylation-regulated kinase 1A
EE: encapsulation efficiency
ENTPD3: ectonucleoside triphosphate di-phosphohydrolase 3
Ex4: Exendin-4
FACS: fluorescence activated cell sorting
GFP: green fluorescent protein
GLP-1: glucagon-like peptide 1
GLP-1R: glucagon-like peptide 1 receptor
HA: hyaluronic acid
HPSC: human pluripotent stem cell
NC: nanocapsule
NC+: nanocapsule positive
NC-: nanocapsule negative
NOD-scid: non-obese diabetic/severe combined immunodeficiency
PBS: phosphate buffered saline
PCL: polycaprolactone
PDI: polydispersity index
PTM: pentamidine
Rh123: rhodamine 123
sBC: cell-derived β-like cells
SEM: scanning electron microscopy
T1D: type 1 diabetes
TEM: transmission electron microscopy
TMR: tetramethylrhodamine
W/O: water in oil
W/O/W: water in oil in water.

## REFERENCES

(1) Gregory, G. A.; Robinson, T. I. G.; Linklater, S. E.; Wang, F.; Colagiuri, S.; de Beaufort, C.; Donaghue, K. C.; Magliano, D. J.; Maniam, J.; Orchard, T. J.; Rai, P.; Ogle, G. D. Global Incidence, Prevalence, and Mortality of Type 1 Diabetes in 2021 with Projection to 2040: A Modelling Study. Lancet Diabetes Endocrinol 2022, 10, 741–760. 10.1016/S2213-8587(22)00218-2.

(2) Paschou, S. A.; Papadopoulou-Marketou, N.; Chrousos, G. P.; Kanaka-Gantenbein, C. On Type 1 Diabetes Mellitus Pathogenesis. Endocr Connect 2018, 7 (1), R38–R46. 10.1530/EC-17-0347.

(3) Merchant, H. J.; McNeilly, A. D. Hypoglycaemia: Still the Main Drawback of Insulin 100 Years on: “From Man to Mouse.” Diabetic Medicine 2021, 38 (12), 1–11. 10.1111/dme.14721.

(4) Wang, P.; Karakose, E.; Choleva, L.; Kumar, K.; De-Vita, R. J.; Garcia-Ocaña, A.; Stewart, A. F. Human Beta Cell Regenerative Drug Therapy for Diabetes: Past Achievements and Future Challenges. Front Endocrinol (Lausanne) 2021, 12 (July), 1–20. 10.3389/fendo.2021.671946.

(5) Wang, P.; Karakose, E.; Choleva, L.; Kumar, K.; De-Vita, R. J.; Garcia-Ocaña, A.; Stewart, A. F. Human Beta Cell Regenerative Drug Therapy for Diabetes: Past Achievements and Future Challenges. Front Endocrinol (Lausanne) 2021, 12 (July), 1–20. 10.3389/fendo.2021.671946.

(6) Tahtouh, T.; Elkins, J. M.; Filippakopoulos, P.; Soundararajan, M.; Burgy, G.; Durieu, E.; Cochet, C.; Schmid, R. S.; Lo, D. C.; Delhommel, F.; Oberholzer, A. E.; Pearl, L. H.; Carreaux, F.; Bazureau, J. P.; Knapp, S.; Meijer, L. Selectivity, Cocrystal Structures, and Neuroprotective Properties of Leucettines, a Family of Protein Kinase Inhibitors Derived from the Marine Sponge Alkaloid Leucettamine B. J Med Chem 2012, 55 (21), 9312–9330. 10.1021/jm301034u.

(7) Ovalle, F.; Grimes, T.; Xu, G.; Patel, A. J.; Grayson, T. B.; Thielen, L. A.; Li, P.; Shalev, A. Verapamil and Beta Cell Function in Adults with Recent-Onset Type Diabetes. Nat Med 2018, 24, 1108–1112. 10.1038/s41591-018-0089-4.

(8) Chenthamara, D.; Subramaniam, S.; Ramakrishnan, S. G.; Krishnaswamy, S.; Essa, M. M.; Lin, F. H.; Qoronfleh, M. W. Therapeutic Efficacy of Nanoparticles and Routes of Administration. Biomater Res 2019, 23 (1). 10.1186/s40824-019-0166-x.

(9) Fillat, C.; Jose, A.; Ros, X. B. De; Mato-Berciano, A.; Maliandi, M. V.; Sobrevals, L. Pancreatic Cancer Gene Therapy: From Molecular Targets to Delivery Systems. Cancers (Basel) 2011, 3, 368–395. 10.3390/cancers3010368.

(10) Blanco, E.; Shen, H.; Ferrari, M. Principles of Nanoparticle Design for Overcoming Biological Barriers to Drug Delivery. Nat Biotechnol 2015, 33 (9), 941–951. 10.1038/nbt.3330.

(11) Kim, K.; Choi, H.; Choi, E. S.; Park, M. H.; Ryu, J. H. Hyaluronic Acid-Coated Nanomedicine for Targeted Cancer Therapy. Pharmaceutics 2019, 11 (7), 1–22. 10.3390/pharmaceutics11070301.

(12) Lu, Q.; Yu, H.; Zhao, T.; Zhu, G.; Li, X. Nanoparticles with Transformable Physicochemical Properties for Overcoming Biological Barriers. Nanoscale. Royal Society of Chemistry July 14, 2023, pp 13202–13223. 10.1039/d3nr01332d.

(13) De, M.; Ghosh, P. S.; Rotello, V. M. Applications of Nanoparticles in Biology. Advanced Materials 2008, 20 (22), 4225–4241. 10.1002/adma.200703183.

(14) Quintanar-Guerrero, D.; Allémann, E.; Fessi, H.; Doelker, E. Preparation Techniques and Mechanisms of Formation of Biodegradable Nanoparticles from Preformed Polymers. Drug Dev Ind Pharm 1998, 24 (12), 1113–1128. 10.3109/03639049809108571.

(15) Lee, C.; Li, Y.; Huang, C.; Lai, J. Poly(ε-Caprolactone) Nanocapsule Carriers with Sustained Drug Release: Single Dose for Long-Term Glaucoma Treatment. Nanoscale 2017, 9, 11754–11764. 10.1039/c7nr03221h.

(16) Memisoglu-Bilensoy, E.; Şen, M.; Hincal, A. A. Effect of Drug Physicochemical Properties on in Vitro Characteristics of Amphiphilic Cyclodextrin Nanospheres and Nanocapsules. J Microencapsul 2006, 23 (1), 59– 68. 10.1080/02652040500286227.

(17) Mondal, D.; Griffith, M.; Venkatraman, S. S. Polycaprolactone-Based Biomaterials for Tissue Engineering and Drug Delivery: Current Scenario and Challenges. International Journal of Polymeric Materials and Polymeric Biomaterials 2016, 65 (5), 255–265. 10.1080/00914037.2015.1103241.

(18) Bartnikowski, M.; Dargaville, T. R.; Ivanovski, S.; Hutmacher, D. W. Degradation Mechanisms of Polycaprolactone in the Context of Chemistry, Geometry and Environment. Prog Polym Sci 2019, 96, 1–20. 10.1016/j.progpolymsci.2019.05.004.

(19) Marchal-Heussler, L.; Sirbat, D.; Hoffman, M.; Maincent, P. Poly(ε-Caprolactone) Nanocapsules in Carteolol Ophthalmic Delivery. Pharmaceutical Research: An Official Journal of the American Association of Pharmaceutical Scientists. 1993, pp 386–390. 10.1023/A:1018936205485.

(20) Woodruff, M. A.; Hutmacher, D. W. The Return of a Forgotten Polymer - Polycaprolactone in the 21st Century. Progress in Polymer Science (Oxford) 2010, 35 (10), 1217–1256. 10.1016/j.progpolymsci.2010.04.002.

(21) Palamà, I. E.; Cortese, B.; D’Amone, S.; Gigli, G. MRNA Delivery Using Non-Viral PCL Nanoparticles. Biomater Sci 2015, 3 (1), 144–151. 10.1039/c4bm00242c.

(22) Ubrich, N.; Bouillot, P.; Pellerin, C.; Hoffman, M.; Maincent, P. Preparation and Characterization of Propranolol Hydrochloride Nanoparticles: A Comparative Study. Journal of Controlled Release 2004, 97 (2), 291–300. 10.1016/j.jconrel.2004.03.023.

(23) Mustafa, G.; Hassan, D.; Ruiz-Pulido, G.; Pourmadadi, M.; Eshaghi, M. M.; Behzadmehr, R.; Tehrani, F. S.; Rahdar, A.; Medina, D. I.; Pandey, S. Nanoscale Drug Delivery Systems for Cancer Therapy Using Paclitaxel— A Review of Challenges and Latest Progressions. Journal of Drug Delivery Science and Technology. Editions de Sante June 1, 2023. 10.1016/j.jddst.2023.104494.

(24) Nezhadi, S.; Saadat, E.; Handali, S.; Dorkoosh, F. Nanomedicine and Chemotherapeutics Drug Delivery: Challenges and Opportunities. J Drug Target 2021, 29 (2), 185–198. 10.1080/1061186X.2020.1808000.

(25) Lee, H.; Ahn, C. H.; Park, T. G. Poly[Lactic-Co-(Glycolic Acid)]-Grafted Hyaluronic Acid Copolymer Micelle Nanoparticles for Target-Specific Delivery of Doxorubicin. Macromol Biosci 2009, 9 (4), 336–342. 10.1002/mabi.200800229.

(26) Babič, A.; Vinet, L.; Chellakudam, V.; Janikowska, K.; Allémann, E.; Lange, N. Squalene-PEG-Exendin as High-Affinity Constructs for Pancreatic Beta-Cells. Bioconjug Chem 2018, 29 (8), 2531–2540. 10.1021/acs.bioconjchem.8b00186.

(27) Fujita, N.; Fujimoto, H.; Hamamatsu, K.; Murakami, T.; Kimura, H.; Toyoda, K.; Saji, H.; Inagaki, N. Noninvasive Longitudinal Quantification of β-Cell Mass with [111In]-Labeled Exendin-4. FASEB J 2019, 33 (11), 11836–11844. 10.1096/fj.201900555RR.

(28) Jansen, T. J. P.; van Lith, S. A. M.; Boss, M.; Brom, M.; Joosten, L.; Béhé, M.; Buitinga, M.; Gotthardt, M. Exendin-4 Analogs in Insulinoma Theranostics. J Labelled Comp Radiopharm 2019, 62 (10), 656–672. 10.1002/jlcr.3750.

(29) Sowa-Staszczak, A.; Pach, D.; Mikołajczak, R.; Mäcke, H.; Jabrocka-Hybel, A.; Stefańska, A.; Tomaszuk, M.; Janota, B.; Gilis-Januszewska, A.; Małecki, M.; Kamiński, G.; Kowalska, A.; Kulig, J.; Matyja, A.; Osuch, C.; Hubalewska-Dydejczyk, A. Glucagon-like Peptide-1 Receptor Imaging with [Lys40(Ahx-HYNIC-99mTc/EDDA)NH2]-Exendin-4 for the Detection of Insulinoma. Eur J Nucl Med Mol Imaging 2012, 40 (4), 524–531. 10.1007/s00259-012-2299-1.

(30) Yang, B.; Cai, H.; Qin, W.; Zhang, B.; Zhai, C.; Jiang, B.; Wu, Y. Bcl-2-Functionalized Ultrasmall Superparamagnetic Iron Oxide Nanoparticles Coated with Amphiphilic Polymer Enhance the Labeling Efficiency of Islets for Detection by Magnetic Resonance Imaging. Int J Nanomedicine 2013, 8, 3977–3990. 10.2147/IJN.S52058.

(31) Zhang, B.; Yang, B.; Zhai, C.; Jiang, B.; Wu, Y. The Role of Exendin-4-Conjugated Superparamagnetic Iron Oxide Nanoparticles in Beta-Cell-Targeted MRI. Biomaterials 2013, 34 (23), 5843–5852. 10.1016/j.biomaterials.2013.04.021.

(32) Fang, Z.; Chen, S.; Manchanda, Y.; Bitsi, S.; Pickford, P.; David, A.; Shchepinova, M. M.; Corr, I. R.; Hodson, D. J.; Broichhagen, J.; Tate, E. W.; Reimann, F.; Salem, V.; Rutter, G. A.; Tan, T.; Bloom, S. R.; Tomas, A.; Jones, B. Ligand-Specific Factors Influencing GLP-1 Receptor Post-Endocytic Trafficking and Degradation in Pancreatic Beta Cells. 2020, 1–24.

(33) Ast, J.; Novak, A. N.; Podewin, T.; Fine, N. H. F.; Jones, B.; Tomas, A.; Birke, R.; Roßmann, K.; Mathes, B.; Eichhorst, J.; Lehmann, M.; Linnemann, A. K.; Hodson, D. J.; Broichhagen, J. Expanded LUXendin Color Palette for GLP1R Detection and Visualization In Vitro and In Vivo. JACS Au 2022, 2 (4), 1007–1017. 10.1021/jacsau.2c00130.

(34) Jansen, T. J. P.; van Lith, S. A. M.; Boss, M.; Brom, M.; Joosten, L.; Béhé, M.; Buitinga, M.; Gotthardt, M. Exendin-4 Analogs in Insulinoma Theranostics. J Labelled Comp Radiopharm 2019, 62 (10), 656–672. 10.1002/jlcr.3750.

(35) Badran, M. M.; Alomrani, A. H.; Harisa, G. I.; Ashour, A. E.; Kumar, A.; Yassin, A. E. Novel Docetaxel Chitosan-Coated PLGA/PCL Nanoparticles with Magnified Cytotoxicity and Bioavailability. Biomedicine and Pharmacotherapy 2018, 106, 1461–1468. 10.1016/j.biopha.2018.07.102.

(36) Barbault-foucher, S.; Gref, R.; Russo, P.; Guechot, J.; Bochot, A. Design of Poly-ε-Caprolactone Nanospheres Coated with Bioadhesive Hyaluronic Acid for Ocular Delivery. Journal of Controlled Release 2002, 83 (3), 365–375. 10.1016/s0168-3659(02)00207-9.

(37) Barbault-foucher, S.; Gref, R.; Russo, P.; Guechot, J.; Bochot, A. Design of Poly-ε-Caprolactone Nano-spheres Coated with Bioadhesive Hyaluronic Acid for Ocular Delivery. Journal of Controlled Release 2002, 83 (3), 365–375. 10.1016/s0168-3659(02)00207-9.

(38) Safwat, M. A.; Soliman, G. M.; Sayed, D.; Attia, M. A. Gold Nanoparticles Capped with Benzalkonium Chloride and Poly (Ethylene Imine) for Enhanced Loading and Skin Permeability of 5-Fluorouracil. Drug Dev Ind Pharm 2017, 43 (11), 1780–1791. 10.1080/03639045.2017.1339082.

(39) Hassan, B.; Hadi, H. Magnetic Solid-Phase Extraction Based on Benzalkonium Chloride-Coated Fe3O4@SiO2 Nanoparticles for Spectrophotometric Determination of Ritodrine Hydrochloride and Salbutamol Sulfate in Water and Urine Samples. Microchemical Journal 2022, 181. 10.1016/j.microc.2022.107805.

(40) Lee, C.; Li, Y.; Huang, C.; Lai, J. Poly(ε-Caprolactone) Nanocapsule Carriers with Sustained Drug Release: Single Dose for Long-Term Glaucoma Treatment. Nanoscale 2017, 9, 11754–11764. 10.1039/c7nr03221h.

(41) Hernández-Silva, E.; Vázquez-Hernández, F.; Mendoza-Acevedo, S.; Pérez-González, M.; Tomás-Velázquez, S.; Rodríguez-Fragoso, P.; Mendoza-Álvarez, J.; Luna-Arias, J. P. Effect of Stirring Rate on the Size of Hydroxyapatite Nanoparticles Synthesized by a Modified Heat-Treated Precipitation Method. Processing and Application of Ceramics 2023, 17 (2), 133–139. 10.2298/PAC2302133H.

(42) Li, Z. Z.; Wen, L. X.; Shao, L.; Chen, J. F. Fabrication of Porous Hollow Silica Nanoparticles and Their Applications in Drug Release Control. Journal of Controlled Release 2004, 98 (2), 245–254. 10.1016/j.jconrel.2004.04.019.

(43) Jovanovic, A. V.; Underhill, R. S.; Bucholz, T. L.; Duran, R. S. Oil Core and Silica Shell Nanocapsules: Toward Controlling the Size and the Ability To Sequester Hydrophobic Compounds. Chemistry of Materials 2005, 17 (13), 3375–3383. 10.1021/cm0480723.

(44) Areny-Balagueró, A.; Mekseriwattana, W.; Camprubí-Rimblas, M.; Stephany, A.; Roldan, A.; Solé-Porta, A.; Artigas, A.; Closa, D.; Roig, A. Fluorescent PLGA Nanocarriers for Pulmonary Administration: Influence of the Surface Charge. Pharmaceutics 2022, 14. 10.3390/pharmaceutics14071447.

(45) Chansaenpak, K.; Tanjindaprateep, S.; Chaicharoenau-domrung, N.; Weeranantanapan, O.; Noisa, P.; Kamkaew, A. Aza-BODIPY Based Polymeric Nanoparticles for Cancer Cell Imaging. RSC Adv 2018, 8, 39248–39255. 10.1039/c8ra08145j.

(46) Paoletti, F.; Ainger, K.; Donati, I.; Scardigli, R.; Vetere, A.; Cattaneo, A.; Campa, C. Novel Fluorescent Cycloheximide Derivatives for the Imaging of Protein Synthesis. Biochem Biophys Res Commun 2010, 396, 258–264. 10.1016/j.bbrc.2010.04.075.

(47) Tosi, G.; Costantino, L.; Rivasi, F.; Ruozi, B.; Leo, E.; Vergoni, A. V.; Tacchi, R.; Bertolini, A.; Vandelli, M. A.; Forni, F. Targeting the Central Nervous System: In Vivo Experiments with Peptide-Derivatized Nanoparticles Loaded with Loperamide and Rhodamine-123. Journal of Controlled Release 2007, 122 (1), 1–9. 10.1016/j.jconrel.2007.05.022.

(48) Treekoon, J.; Chansaenpak, K.; Tumcharern, G.; Zaiman Zain, Z. S.; Lee, H. B.; Kue, C. S.; Kamkaew, A. Aza-BODIPY Encapsulated Polymeric Nanoparticles as an Effective Nanodelivery System for Photodynamic Cancer Treatment. Mater Chem Front 2021, 5 (5), 2283–2293. 10.1039/d0qm00891e.

(49) Santos, J.; Sousa, F.; Queiroz, J.; Costa, D. Rhodamine Based Plasmid DNA Nanoparticles for Mitochondrial Gene Therapy. Colloids Surf B Biointerfaces 2014, 121, 129–140. 10.1016/j.colsurfb.2014.06.003.

(50) Makama, S.; Kloet, S. K.; Piella, J.; van den Berg, H.; de Ruijter, N. C. A.; Puntes, V. F.; Rietjens, I. M. C. M.; van den Brink, N. W. Effects of Systematic Variation in Size and Surface Coating of Silver Nanoparticles on Their In Vitro Toxicity to Macrophage Raw 264.7 Cells. Toxicological Sciences 2018, 162 (1), 79– 88. 10.1093/toxsci/kfx228.

(51) Abbasi, R.; Shineh, G.; Mobaraki, M.; Doughty, S.; Tayebi, L. Structural Parameters of Nanoparticles Affecting Their Toxicity for Biomedical Applications: A Review; Springer Netherlands, 2023; Vol. 25. 10.1007/s11051-023-05690-w.

(52) Nandhakumar, S.; Dhanaraju, M. D.; Sundar, V. D.; Heera, B. Influence of Surface Charge on the in Vitro Protein Adsorption and Cell Cytotoxicity of Paclitaxel Loaded Poly(ε-Caprolactone) Nanoparticles. Bulletin of Faculty of Pharmacy, Cairo University 2017, 55 (2), 249–258. 10.1016/j.bfopcu.2017.06.003.

(53) Zelenková, T.; Fissore, D.; Marchisio, D. L.; Barresi, A. Size Control in Production and Freeze-Drying of Poly-ε-Caprolactone Nanoparticles. J Pharm Sci 2014, 103, 1839–1850. 10.1002/jps.23960.

(54) Zelenková, T.; Fissore, D.; Marchisio, D. L.; Barresi, A. A. Size Control in Production and Freeze-Drying of Poly-ε-Caprolactone Nanoparticles. J Pharm Sci 2014, 103, 1839–1850. 10.1002/jps.23960.

(55) Selby, L. I.; Cortez-Jugo, C. M.; Such, G. K.; Johnston, P. R. Nanoescapology: Progress toward Understanding the Endosomal Escape of Polymeric Nanoparticles. WIREs Nanomed Nanobiotechnol 2017, 9. 10.1002/wnan.1452.

(56) Docherty, F. M.; Riemondy, K. A.; Castro-gutierrez, R.; Dwulet, J. M.; Shilleh, A. H.; Hansen, M. S.; Williams, S. P. M.; Armitage, L. H.; Santostefano, K. E.; Wallet, M. A.; Mathews, C. E.; Triolo, T. M.; Benninger, R. K. P.; Russ, H. A. ENTPD3 Marks Mature Stem Cell–Derived β-Cells Formed by Self-Aggregation In Vitro. Diabetes 2021, 70 (November), 2554– 2567. 10.2337/db20-0873ISLET.

(57) Saunders, D. C.; Brissova, M.; Phillips, N.; Shrestha, S.; Walker, J. T.; Aramandla, R.; Poffenberger, G.; Flaherty, D. K.; Weller, K. P.; Pelletier, J.; Cooper, T.; Goff, M. T.; Virostko, J.; Shostak, A.; Dean, E. D.; Greiner, D. L.; Shultz, L. D.; Prasad, N.; Levy, S. E.; Carnahan, R. H.; Dai, C.; Sévigny, J.; Powers, A. C. Ectonucleoside Triphosphate Diphosphohydrolase-3 Antibody Targets Adult Human Pancreatic β Cells for In Vitro and In Vivo Analysis. Cell Metab 2019, 29 (3), 745-754.e4. 10.1016/j.cmet.2018.10.007.

(58) Kyosseva, S.V; Harris, E.N.; Weigel, P. H. The Hyaluronan Receptor for Endocytosis Mediates Hyaluronan-Dependent Signal Transduction via Extracellular Signal-Regulated Kinases *. Journal of Biological Chemistry 2008, 283 (22), 15047–15055. 10.1074/jbc.M709921200.

(59) Necker, H. Pentamidine, A New Diabetogenic Drug in Laboratory Rodents P. 1983, 418–423.

(60) Wang, J.; Shi, A.; Agyei, D.; Wang, Q. Formulation of Water-in-Oil-in-Water (W/O/W) Emulsions Containing Trans-Resveratrol. RSC Adv 2017, 7, 35917– 35927. 10.1039/c7ra05945k.

(61) Micallef, S. J.; Li, X.; Schiesser, J. V; Hirst, C. E.; Yu, Q. C.; Lim, S. M.; Nostro, M. C.; Elliott, D. A.; Sarangi, F.; Harrison, L. C.; Keller, G.; Elefanty, A. G.; Stanley, E. G. INSGFP/Whuman Embryonic Stem Cells Facilitate Isolation of in Vitro Derived Insulin-Producing Cells. Diabetologia 2012, 55 (3), 694–706. 10.1007/s00125-011-2379-y.

(62) Docherty, F. M.; Riemondy, K. A.; Castro-Gutierrez, R.; Dwulet, J. M.; Shilleh, A. H.; Hansen, M. S.; Williams, S. P. M.; Armitage, L. H.; Santostefano, K. E.; Wallet, M. A.; Mathews, C. E.; Triolo, T. M.; Benninger, R. K. P.; Russ, H. A. ENTPD3 Marks Mature Stem Cell–Derived β-Cells Formed by Self-Aggregation In Vitro. Diabetes 2021, 70 (11), 2554–2567. 10.2337/db20-0873.

(63) Shilleh, A. H.; Beard, S.; Russ, H. A. Enrichment of Stem Cell-Derived Pancreatic Beta-like Cells and Controlled Graft Size through Pharmacological Removal of Proliferating Cells. Stem Cell Reports 2023, 18 (6), 1284–1294. 10.1016/j.stemcr.2023.05.010.

(64) Yu, Y.; Chau, Y. One-Step “Click” Method for Generating Vinyl Sulfone Groups on Hydroxyl-Containing Water-Soluble Polymers. Biomacromolecules 2012, 13 (3), 937–942. 10.1021/bm2014476.

(65) Podewin, T.; Ast, J.; Broichhagen, J.; Fine, N. H. F.; Nasteska, D.; Leippe, P.; Gailer, M.; Buenaventura, T.; Kanda, N.; Jones, B. J.; M’Kadmi, C.; Baneres, J.-L.; Marie, J.; Tomas, A.; Trauner, D.; Hoffmann-Röder, A.; Hodson, D. J. Conditional and Reversible Activation of Class A and B G Protein-Coupled Receptors Using Tethered Pharmacology. ACS Cent Sci 2018, 4 (2), 166–179. 10.1021/acscentsci.7b00237.

(66) Kong, J.-H.; Oh, E. J.; Chae, S. Y.; Lee, K. C.; Hahn, S. K. Long Acting Hyaluronate – Exendin 4 Conjugate for the Treatment of Type 2 Diabetes. Biomaterials 2010, 31 (14), 4121–4128. 10.1016/j.biomaterials.2010.01.091.

